# Light-induced modulation of DNA recognition by the Rad4/XPC damage sensor protein

**DOI:** 10.1101/2020.09.28.313114

**Authors:** Amirrasoul Tavakoli, Debamita Paul, Hong Mu, Jagannath Kuchlyan, Saroj Baral, Anjum Ansari, Suse Broyde, Jung-Hyun Min

## Abstract

Biomolecular structural changes upon binding/unbinding are key to their functions. However, characterization of such dynamical processes with high spatial and temporal resolutions is difficult as it requires ways to rapidly trigger the assembly/disassembly as well as ways to monitor the structural changes over time. Recently, various chemical strategies have been developed to use light to trigger changes in oligonucleotide structures, thereby their activities. Here we report that photoswitchable DNA can be used to modulate the DNA binding of the Rad4/XPC DNA repair complex using light. Rad4/XPC specifically binds to diverse helix-destabilizing/distorting lesions including bulky organic adducts and functions as a key initiator for the eukaryotic nucleotide excision repair (NER) pathway. We show that the 6-nitropiperonyloxymethyl (NPOM)-modified DNA is recognized by the Rad4 protein as a specific substrate and that the specific binding can be abolished by light-induced cleavage of the NPOM group from DNA in a dose-dependent manner. Fluorescence lifetime-based analyses of the DNA conformations suggest that free NPOM-DNA retains B-DNA-like conformations despite its bulky NPOM adduct, but Rad4-binding renders it to be heterogeneously distorted. Subsequent extensive conformational searches and molecular dynamics simulations demonstrate that NPOM in DNA can be housed in the major groove of the DNA, with stacking interactions among the nucleotide pairs remaining largely unperturbed and thus retaining overall B-DNA conformation. Our work suggests that photoactivable DNA can be used as a DNA lesion surrogate to study DNA repair mechanisms such as nucleotide excision repair.

Graphical Summary
This work shows that a photolabile 6-nitropiperonyloxymethyl (NPOM)-modified DNA is specifically recognized by the Rad4/XPC damage sensor protein complex that initiates the nucleotide excision repair pathway; light-induced cleavage of NPOM abolishes the specific binding to Rad4/XPC.

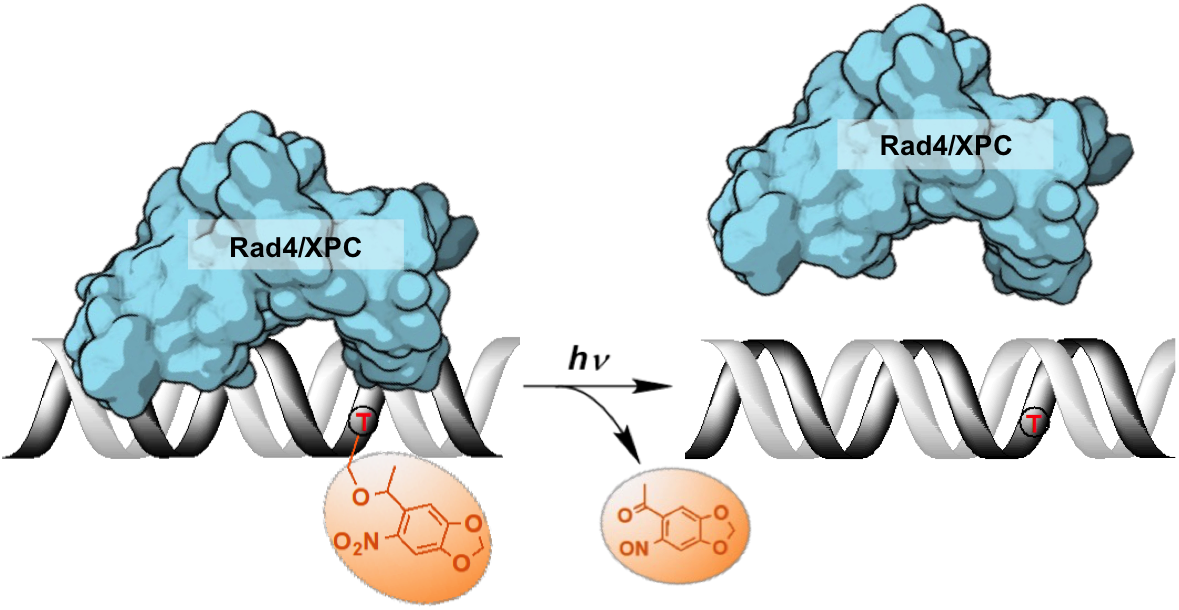

## INTRODUCTION

Biological processes entail dynamic yet coordinated assembly and disassembly of multiple molecules in solution. A key challenge in studying these processes in high structural and temporal resolution lies in the difficulties in controlled triggering of these events. Methods of triggers often involve either perturbing the equilibrium state using temperature-, salt- or pH-jumps or initiating the binding/unbinding through rapid mixing. Another method of triggering involves photo-induced chemical changes, as showcased by pioneering studies on dynamics of allosteric transitions in hemoglobin^1–2^. Photoswitchable groups undergo structural changes upon irradiation by light, usually of a specific wavelength, in a reversible or irreversible manner^3–4^ Such light-induced chemical/structural conversions have emerged as useful tools to control the properties and hence probe the functions of biomolecules harboring the photoswitchable groups, as light can be easily applied to biological systems in vitro or in vivo to trigger specific events^5–9^. Photoswitchable modifications in small molecules^10–12^, oligonucleotides^13^, peptides^14–15^, and proteins (mostly enzymes)^16–19^ have also been used to control and monitor a wide variety of biological outcomes including gene expression, enzyme activity, oligomerization states, cellular localization and immune responses.

One of the photoswitchable groups, the 6-nitropiperonyloxymethyl (NPOM) has been introduced by the Deiters group as an improvement over previous *o*-nitrobenzyl derivatives for modifying biomolecules including oligonucleotides^20–22^ (**Figure 1**). Irradiating an NPOM-modified nucleoside/ DNA/RNA with light (λ = 365 nm) cleaves the NPOM from the substrate without damaging the parent compound and restores the unmodified structures with concomitant release of 1-(6-nitroso-1,3-benzodioxol-5-yl)ethenone (hereafter nitrosoacetophenone) (**Figure 1**)^23–26^. Compared with the previous O-nitrobenzyl derivatives such as 6-nitroveratryl (NV) and 6-nitroveratryloxycarbonyl (NVOC) groups, NPOM features a higher quantum yield (Φ = 0.094 versus Φ = 0.0013 for NVOC), is highly stable in an aqueous environment at various pHs^21, 23, 27^ and penetrates cell membranes without altering growth rate or phenotype of the cells/organisms^28^. The NPOM modification has been applied to various *in vitro* and *in vivo* biological studies. For instance, photocleavage of NPOM-modified DNA/RNA has been used to control the activities of DNAzymes^29^, antisense DNA/RNA^13, 30^, restriction endonucleases^31^, DNA-binding transcription factors^32^, polymerase chain reaction (PCR) rates^33^, as well as CRISPR-Cas gene editing^9, 18, 34–35^. In these cases, NPOM-modifications prevent the DNA or RNA from hybridizing to the complementary strands, which could be reversed upon NPOM cleavage with light, converting the unhybridized ‘inactive’ molecules to hybridized ‘active’ molecules.

**Figure 1.**
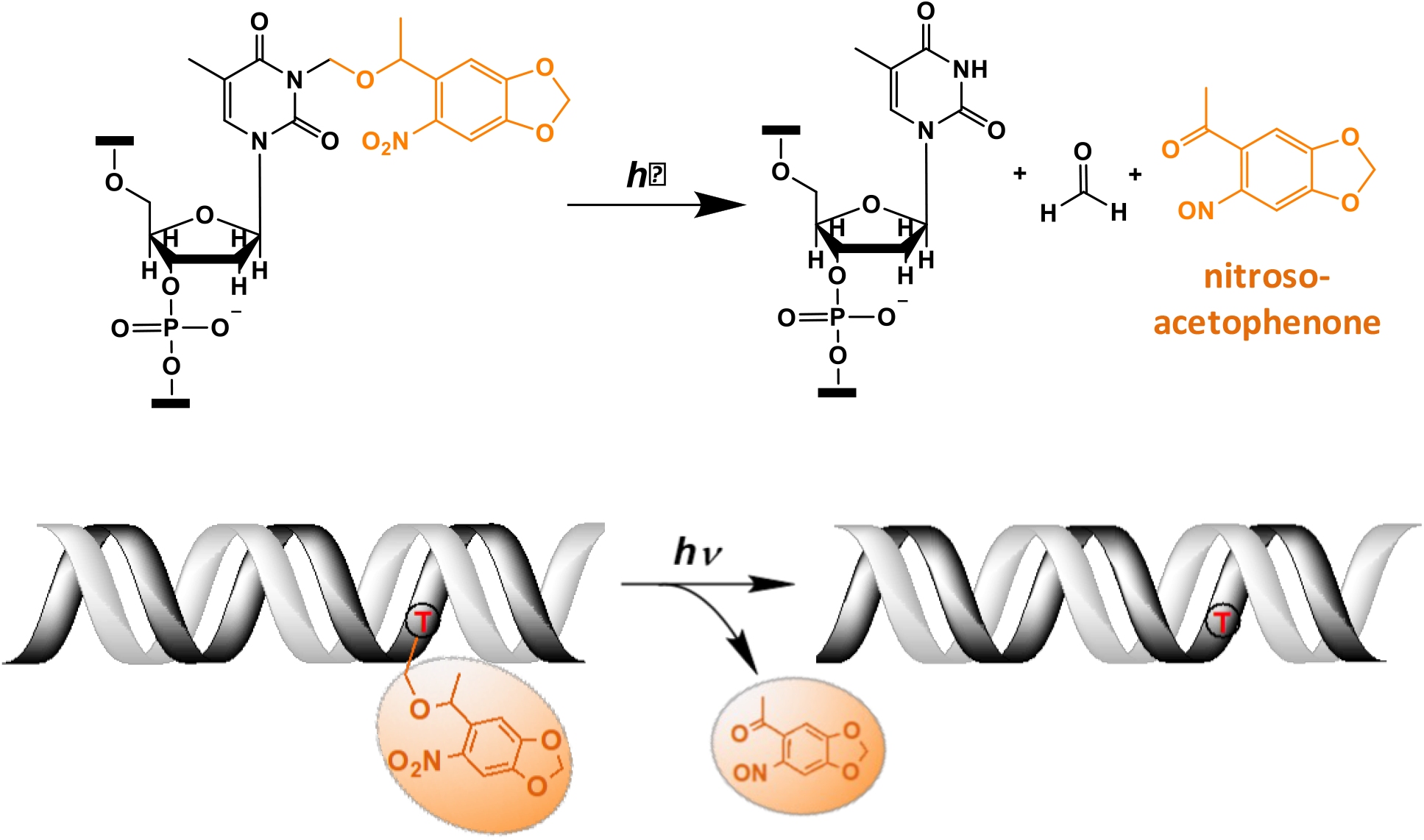
Light-induced photocleavage reaction of the NPOM group from DNA. (*Top*) Schematic of the photocleavage reaction. Upon light irradiation (λ=365nm), the NPOM group (orange) is cleaved from the modified thymidine (dT) in the DNA, restoring unmodified thymidine while releasing nitrosoacetophenone. (*Bottom*) Cartoon of the photocleavage reaction in NPOM-containing duplexed DNA.

Here, we took advantage of the fact that DNA modifications such as NPOM are quite bulky and thus perhaps could be seen as a lesion on DNA by cellular DNA repair machineries, particularly those involved in the nucleotide excision repair (NER) pathway. NER repairs a wide spectrum of bulky adduct lesions in the DNA including sunlight-induced intra-strand crosslinks, bulky DNA adducts induced by various metabolites, reactive oxygen species, environmental pollutants and carcinogens (reviewed in^36–38^). Genetic impairment in NER causes high sun sensitivity xeroderma pigmentosum (XP) cancer predisposition syndrome in humans^37, 39^. In eukaryotic NER, the repair of these lesions scattered around the global genome is primarily initiated when the XPC–RAD23B complex (Rad4–Rad23 in yeast; hereafter referred to as XPC and Rad4) first specifically localizes to the lesion. The lesion binding by Rad4/XPC subsequently leads to the recruitment to the lesion of the transcription factor IIH complex (TFIIH) containing XPD and XPB helicases, which verifies the presence of a bulky lesion and recruits other NER factors. Eventually, a 24–32 nucleotide (nt) lesion-containing portion of the DNA strand is excised by the XPF-ERCC1 and XPG endonucleases and the gap in the DNA is restored by repair synthesis and nick sealing.

Previous studies from our group have shown that Rad4 recognizes DNA lesions in an indirect manner: crystal structures of Rad4 bound to UV-lesions showed that the Rad4 flips two damage-containing nucleotide pairs out of the duplex (the so-called ‘open’ conformation) with the damaged nucleotides flipped away from the protein, such that only the undamaged nucleotides on the complementary strand make direct contacts with the protein^40–41^. This and other studies pose a puzzle as to the mechanism of this indirect readout by Rad4 and the nature of the structural intermediates along the recognition trajectory. These missing key steps led us to ponder whether photoswitchable adducts could serve as model DNA lesions and if their photoswitchable characteristics could be used to trigger the binding/unbinding events in a precisely controlled manner for structural and functional studies. So far, photoswitchable DNA/RNA has been mostly used for cellular and genetic studies but not as much for probing the biochemical and structural mechanisms, let alone for investigating NER^9, 42^.

Here, using fluorescence lifetime-based fluorescence resonance energy transfer (FRET) measurements with cytosine analog FRET pair, tC° and tC_nitro_, that are uniquely sensitive to changes in B-DNA conformation^43–46^, together with UV-Vis spectroscopy and competitive electrophoretic mobility shift assays (EMSA), we show that (1) the NPOM modification on DNA is recognized specifically by Rad4 and (2) that this specific binding is completely abolished by photocleavage of the NPOM moiety from the DNA. Extensive conformational searches and molecular dynamics simulations of the DNA also corroborate with the fluorescence-based conformational analyses; the results demonstrate that NPOM in DNA can be housed in the major groove of the DNA, with stacking interactions among the nucleotide pairs remaining largely unperturbed and thus retaining overall B-DNA conformation. These findings pave the way for the use of photoswitchable DNA as a novel probe to examine the damage recognition mechanism in the NER pathway and is a significant addition to the toolbox to study the structure and dynamics of molecules.

## MATERIALS AND METHODS

### Preparation of Rad4–Rad23 complexes

The Rad4-Rad23 complex (or simply referred to as “Rad4”) was prepared as previously described^40–41, 47^ The Rad4 construct spans residues 101–632 and contains all four domains involved in DNA binding. This Rad4-Rad23 construct has previously been shown to exhibit the same DNA-binding characteristics as the full-length complex^41^. While Rad23 does not participate in DNA binding directly, it is required for stabilizing Rad4.

Hi5 insect cells co-expressing Rad4 and Rad23 proteins were harvested 2 days after infection. After lysis, the protein complex was purified by affinity chromatography (Ni-NTA Agarose, MCLAB), anion-exchange (Source Q, GE healthcare) and cation exchange (Source S, GE healthcare) chromatography followed by gel filtration (Superdex200, GE healthcare). The chromatogram and SDS-PAGE analyses of the gel filtration step show that peak fractions contain a homogeneous 1:1 complex of Rad4 and Rad23 proteins. These peak fractions were pooled and further concentrated by ultrafiltration (Amicon Ultra-15, Millipore) to ~13–14 mg/ml (135–150 μM) in 5 mM bis-tris propane–HCl (BTP-HCl), 800 mM sodium chloride (NaCl) and 5 mM dithiothreitol (DTT), pH 6.8. The complex was prepared without thrombin digestion, thus retaining the UBL domain of Rad23 and a histidine-tag on Rad4.

### Preparation of double-stranded DNA substrates

Unmodified DNA oligonucleotides were purchased from Biosynthesis or Integrated DNA Technologies (IDT). DNA oligomers synthesized with tC^o^ and tC_nitro_ were from Biosynthesis. All oligonucleotides were purchased as HPLC-purified. Oligonucleotides appeared as a single band on denaturing polyacrylamide gels, indicating high length-purity (>90%) of the oligonucleotides. The concentrations of each singlestranded DNA were determined by UV absorbance at 260 nm (NanoDrop One^C^, ThermoFisher) using the A_260_ extinction coefficients calculated by the nearest neighbor method (Biosynthesis). To prepare DNA duplexes, two complementary oligonucleotides were mixed at 1:1 molar ratio at 100 μM in TE buffer (10 mM Tris-HCl pH 7.50, 1 mM EDTA) in a microcentrifuge tube and annealed by slow-cooling: the tube was immersed in a 1.2 L hot water bath (~100 °C) placed on a hot plate; after 5 minutes, the hot plate was turned off and the samples were cooled to room temperature over 5 to 6 hours.

### Photo-irradiation experiments and cleavage studies with NPOM DNA and NPOM-dT (NPOM-modified deoxythymidine)

NPOM-dT was purchased from Berry & Associates (Cat No. PY 7795). 50 μl of 10-50 μM of DNA samples or NPOM-dT were irradiated in a chamber encasing four UV-A lamps, at 6.4 mW/cm^2^ (measured by General Tools Digital UVA/UVB Meter, 280-400 nm; #UV513AB). At specified time intervals, 1.5 μl of the sample was taken out and its absorbance spectrum was measured with NanoDrop One UV-Vis Spectrophotometer. The experiments were done with lights off in the lab. Exposing the NPOM-dT 10-400 μM in TE buffer with 0.01-0.4% DMSO or NPOM-DNA 10-50 μM in TE buffer under ambient light in the lab for 24 hours did not change the absorbance spectra of the samples, indicating little photocleavage by ambient light.

### Melting temperature measurements of DNA duplexes

The overall thermal stabilities of all DNA duplexes were measured as follows. The absorbance at 260 nm of each DNA duplex (1.5 μM) was measured in a sample cuvette of path length 1 cm, using Cary 300 Bio UV-Visible spectrophotometer equipped with a Varian temperature controller. The absorbance measurements were done from 10 to 85 °C at every 1.0 °C interval. Derivative method^48^ of Carry300 (Thermal software) was used to calculate the melting temperature (T_m_) at which 50% of the DNA strands have separated. The derivatives were obtained numerically from the absorbance data using a Savitzky Golay technique where the difference between adjacent points was first computed followed by a smoothing procedure where 5 points surrounding an individual point were averaged to produce a new, smoothed point^49^.

### Competition electrophoretic mobility shift assays (EMSA)

To determine the relative affinities of Rad4 binding to different DNA substrates, competition EMSA (or gel-shift assays) were employed essentially as previously described^40–41, 43, 47, 50^. The benefit of using this competition assay over the conventional single-substrate EMSA is that one can directly observe any preferential binding over the nonspecific binding, including factors such as potential DNA end-binding while avoiding multiple proteins aggregating on a single DNA, as is the case when protein is in excess of total DNA^50^. Various concentrations of the Rad4-Rad23 complexes were mixed with 5 nM ^32^P-labelled DNA of interest (mismatched/damaged or matched/undamaged) in the presence of 1000 nM cold (unlabeled), matched DNA, CH7_NX in an EMSA buffer (5 mM BTP-HCl, 75 mM NaCl, 5 mM DTT, 5% glycerol, 0.74 mM 3-[(3-cholamidopropyl)dimethylammonio]-1-propanesulfonate (CHAPS), 500 μg ml^-1^ bovine serum albumin, pH 6.8). Mixed samples were incubated at room temperature for 20 min and separated on 4.8% non-denaturing polyacrylamide gels in 1x TBE buffer (89 mM Tris-HCl, 89 mM Boric acid, 2 mM EDTA, pH 8.0), run at constant 150 V for 15 min at 4 °C. The gels were quantitated by autoradiography using Typhoon FLA9000 and Imagelab 6.0.1 software from Biorad. The averages of the Rad4-bound DNA fractions quantified from three independent EMSA gels were used for subsequent calculations of the apparent dissociation constants (K_d,app_).

To obtain apparent dissociation constants (K_d,app_) for different DNA substrates, we first used the matched CH7_NX DNA as both the ‘hot’ probe and the cold competitor DNA, and obtained the K_d,app_ for CH7_NX (K_ns_) by fitting the fraction of labelled DNA bound (f) to the equation

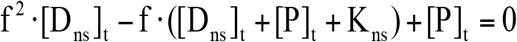

where [P]_t_ is the total protein concentration and [D_ns_]_t_ is the total CH7_NX concentration (1005 nM). The K_d,app_’s of other DNA substrates (K_s_) were subsequently obtained by using the DNA of interest as the ^32^P-labelled DNA probe and fitting the fraction of labelled DNA bound (f) to the equation:

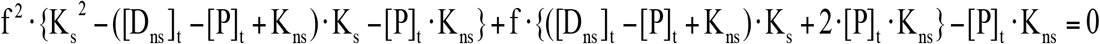

where [P]_t_ is the total protein concentration, [D_ns_]_t_ is the concentration of the undamaged competitor CH7_NX (1000 nM), and K_ns_ is the K_d,app_ for CH7_NX binding, as obtained above. The equation for K_s_ was obtained using the approximation that the concentration of Rad4-bound labelled DNA is negligible compared to the total concentrations of Rad4 and of the matched/undamaged DNA competitor. Curve fittings for K_ns_ and K_s_ were both done by the nonlinear regression method using Origin software (OriginLab). The errors reported for K_d,app_ indicate the errors of the nonlinear regression fit^40–41^.

### Fluorescence lifetime measurements

DNA duplexes labeled with both tC^o^ and tC_nitro_ (DNA_DA) or tC^o^ alone (DNA_D) were prepared as described above. The DNA and Rad4-Rad23-DNA 1:1 complexes were prepared at 5 μM in phosphate-buffered saline (10 mM Na_2_HPO_4_, 2 mM KH_2_PO_4_, 137 mM NaCl, 2.7 mM KCl pH 7.4) with 1 mM DTT. Under this condition, native gel electrophoresis and dynamic light scattering experiments showed that the Rad4-Rad23–DNA samples form uniformly sized 1:1 protein:DNA complexes^50^. Sample volume for each fluorescence lifetime (FLT) measurement was 12 μl. Time-resolved fluorescence decays were measured on a time-correlated single-photon counting (TCSPC) system (DeltaFlex, HORIBA). In our experimental setup, the excitation pulse was pulse-picked second harmonic of a mode-locked Ti-sapphire laser (Mai Tai HP, Spectra-Physics) with a tunable range from 690 to 1040 nm. We used ~100 fs pulses centered at 730 nm with a repetition rate of 80 MHz and average power 1.8 W (~23 nJ/pulse). The fundamental beam (730 nm) was passed through a half-wave plate and then to a pulse picker for reducing the repetition rate of the beam from 80 MHz to 4 MHz. For the second harmonic generation, the fundamental 730 nm beam coming from the pulse picker was focused onto a thin β-barium borate (BBO) crystal in second-harmonic generator TPH Tripler (Minioptics, Inc., Arcadia, CA), which generates a frequency-doubled beam centered at 365 nm. The remaining fundamental beam and frequency-doubled 365 nm beam were separated through a dichroic mirror. The fundamental pulse was given a time delay through an optical delay cable connected with a delay box, and was allowed to pass through a photodiode for triggering the start signal in DeltaFlex collection system. For excitation of tC^o^, the frequency-doubled laser pulses were passed through a monochromator set at 365 nm (band pass 10 nm) attached to the DeltaFlex set up. We used neutral density filter (FSQ-ND20, broadband UV-grade fused silica metallic filter, Newport corporation) to reduce power of excitation light suitable for TCSPC measurements. The intensity of the laser delivered to the samples was 0.21 mW/cm^2^ as measured by a General Tools Digital UVA/UVB Meter, 280-400 nm (#UV513AB).

The fluorescence from the samples passed through a long pass filter cut at 375 nm and the signal was collected at magic angle with the emission polarizer oriented 54.7 from the vertical. The transmitted signal was passed through the entrance slit of emission monochromator set at 470 nm (band pass 10 nm) and collected by a Picosecond Photon Detection module (PPD-850, Horiba) connected to a photon counter. The instrument response function (IRF) of the system was measured using a dilute aqueous solution (3% w/w) of LUDOX AM colloidal silica (Sigma-Aldrich). The full-width half maxima (FWHM) of the instrument response function (IRF) was ~425 ps. Fluorescence decay curves were recorded on a 100 ns timescale, resolved into 4,096 channels, to a total of 10,000 counts in the peak channel. For temperature scanning, the temperature of the sample chamber was regulated by Quantum Northwest Peltier-Controlled TC1, between 4–40°C.

### Analysis of the fluorescence decay traces using discrete exponential (DE) fits

The discrete exponential (DE) analysis was carried out using EzTime software (version 3.2.9.9) provided by Horiba that uses a standard iterative reconvolution method, assuming a multiexponential decay function, 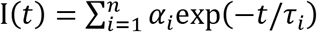, where *α_i_* is the amplitude and *τ_i_* is the fluorescence lifetime of the i-th decay component (**Table S1**). The maximum number of exponentials allowed by this software is five. For all measured decay traces, no more than four exponentials were needed to reasonably fit the data. The number of exponentials required for each trace was determined by the quality of the fit, evaluated based on the reduced chi-square *χ*^2^ and the randomness of residuals (**Figure S4**). Each exponential component for the donor-acceptor labeled samples (DNA_DA) was characterized in terms of a lifetime denoted as *τ_DA,i_*, and a corresponding normalized amplitude or relative population 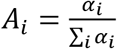. The FRET efficiency for the population in that component was computed from 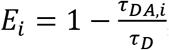, where *τ_D_* indicates the intrinsic lifetime of the donor probe. The average FRET efficiency for each sample was computed as 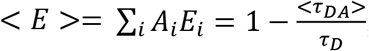, where 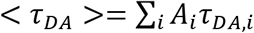. For cases where the intrinsic lifetime of the donor-only samples could not be described by a single exponential, *τ_D_* was taken as the intrinsic lifetime of the donor probe obtained from unmodified DNA (AT10_D).

### Analysis of the decay traces using maximum entropy method (MEM) and Gaussian fitting

Though the DE fitting has been traditionally used for fluorescence lifetime decay analyses, MEM has distinct advantages^43, 51^. In our previous studies, we have also shown that the results from the DE and MEM analyses corroborate with each another^43^. The MEM analyses were carried out using MemExp software^52–53^, as done previously^43^. The reproducibility of the distributions obtained from the MEM analyses from three independent lifetime measurements on each sample are illustrated in (**Figure S6**). The data presented in (**Figure 4**) are for one representative from this set. To further characterize the lifetime distributions from the MEM analyses, we fitted the measured distributions to a sum of Gaussians (**Figure S5**). Each Gaussian component was used to calculate the average FRET representing that component, and the area under the Gaussian curve was taken as a measure of the fractional population of that component. The results are summarized in **Table S2**. Errors are indicated with standard deviations (s.d.) from three independent sets of measurements.

### Conformational searches and molecular dynamics simulations of NPOM-dT-containing DNA duplex structures

In order to explore the structures of the NPOM-dT-containing DNA duplex, we first modeled NPOM-dT at the center of a 13-mer B-DNA duplex with the same sequence as in the AT2 NPOM-DNA (**Table 1**) employed in the experimental study. We carried out extensive conformational searches beginning with an NPOM-dT nucleoside to generate initial models for MD simulations of NPOM-dT in a B-DNA duplex, utilizing a sequence of protocols involving molecular modeling (Discovery Studio 2.5, Accelrys Software Inc.) and quantum mechanical geometry optimization (Gaussian 09^54^) to define sterically feasible NPOM-dT rotamer combinations for initiating the MD simulations (Figures S11-S12). These protocols and obtained structures are summarized in **Scheme 1**. We used the AMBER18 suite of programs^55^ for MD simulations and analyses. Full details of NPOM-dT force field parameterization, MD simulation methods and analyses are given in the SI Methods. Newly developed force field parameters for the NPOM-dT are given in **Table S3**.

**Table 1.**
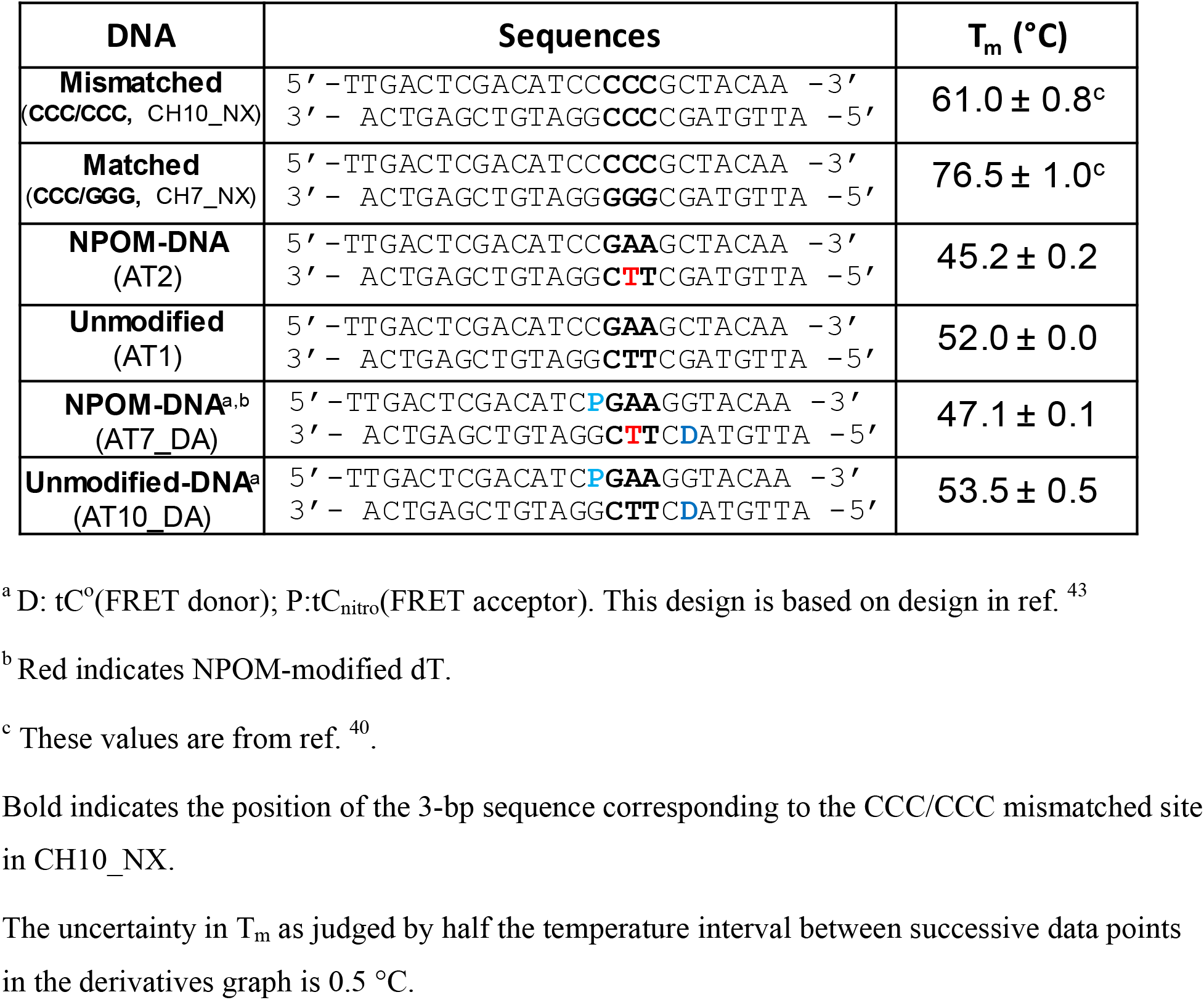
Sequences of the DNA duplexes used in this study.

**Scheme 1.**
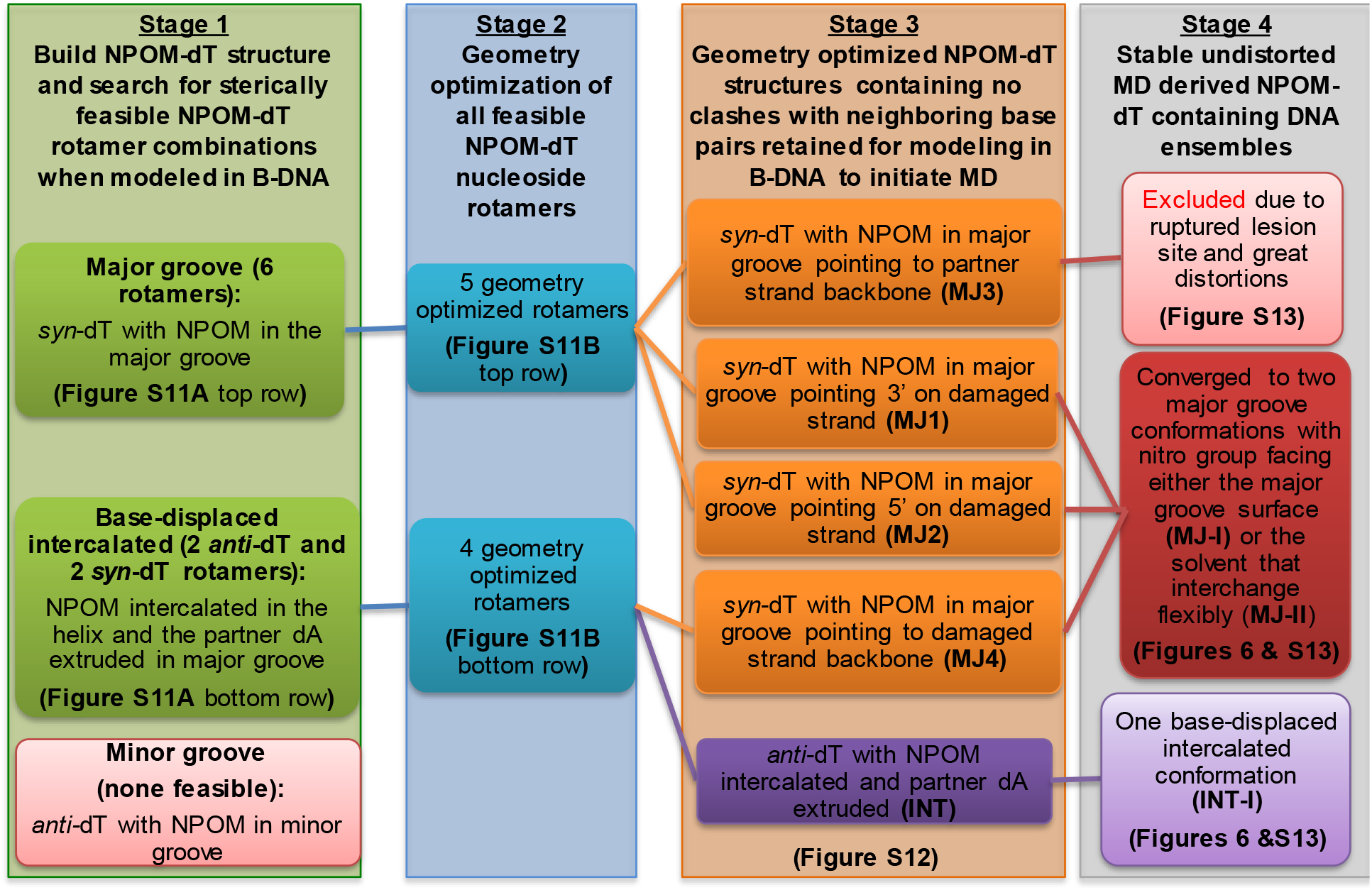
Conformational search strategy for NPOM-dT-containing DNA

## RESULTS

### Dose-dependent photocleavage of NPOM from DNA as monitored by UV-Vis absorption spectroscopy

Photoconversion reactions often induce changes in the absorption spectra of the chemical groups of interest, which in turn can be used to track the reaction progress. To monitor the NPOM’s photocleavage reaction, we obtained the UV-Vis absorption spectra of the NPOM-modified DNA duplex (NPOM-DNA or AT2; see **Table 1** for the DNA sequences used in the study) and dT nucleoside (NPOM-dT) after they were irradiated with varying doses of photocleavage-inducing light (λ=365 nm, 6.4 mW/cm^2^, irradiated for 0 to 210 sec). The overall absorbance in the 300-500 nm range increased with increased irradiation time, with the absorption maximum (*λ_max_*) shifting from 365 nm to 395 nm (red-shift) for both samples (**Figures 2 and S1**). NPOM-DNA and NPOM-dT also showed similar reaction kinetics with similar half-times (t_1/2_) of the absorption change at 395 nm (57 ± 7 sec for NPOM-DNA and 54 ± 5 sec for NPOM-dT) and a common ~3-fold increase in the absorbance upon saturation (**Figure 2A, B**). Unmodified DNA duplex did not show absorption in this wavelength range with or without irradiation (**Figure 2B**). The strong absorbance at 395 nm after photo-irradiation comes from the photocleavage reaction product, nitrosoacetophenone (**Figure 1**)^56^. When the nitroso-acetophenone was removed from DNA using a size-exclusion purification (G25, MWCO ~5 kDa), the absorption spectrum of the sample largely returned to that of the unmodified DNA (**Figure S1D**). Altogether, the results confirm the photo-induced cleavage of NPOM from DNA and indicate that the photocleavage reaction can be modulated by light doses^9^. The photocleavage reaction may, in principle, be accelerated by using light of higher intensity and shorter time duration, as indicated previously^9, 57^ Although there have been multiple studies using NPOM as photoactivatable group that elicit various biological outcomes, this is the first time the photocleavage reaction progress was characterized *in situ* (through monitoring of the absorption spectra).

**Figure 2.**
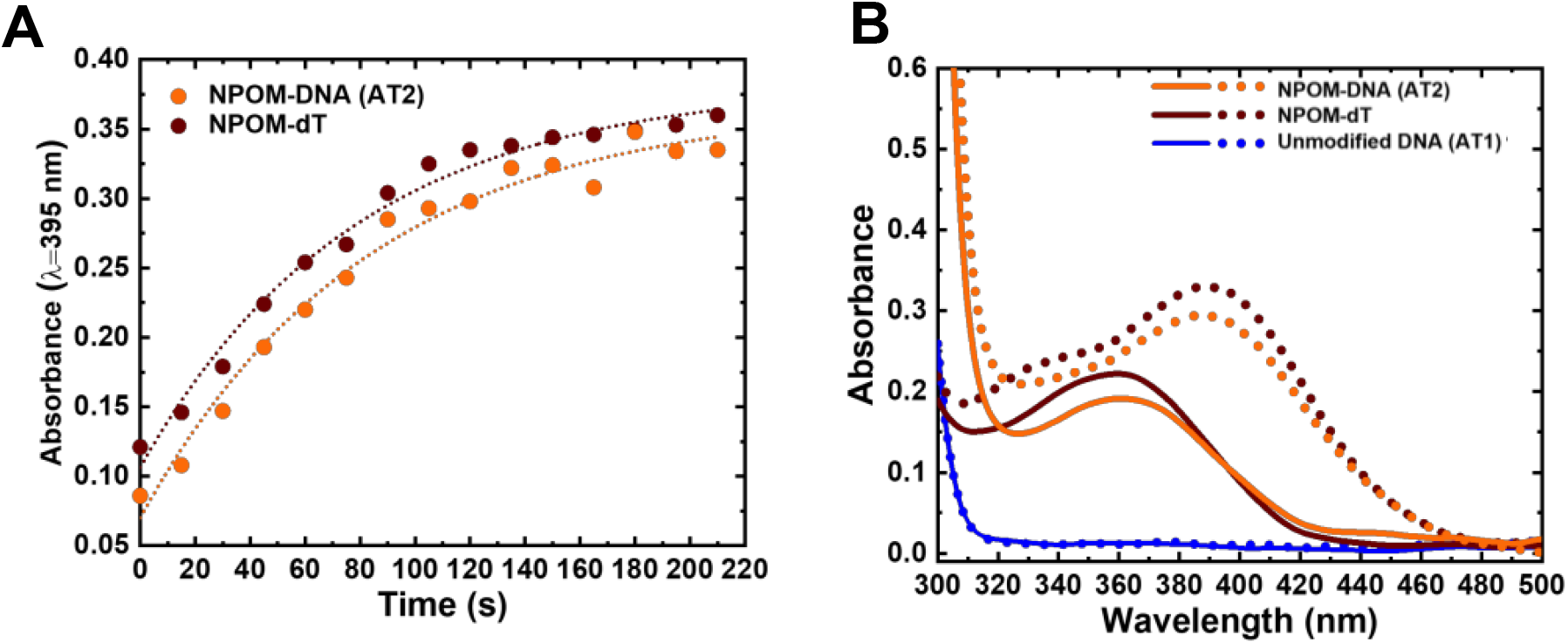
Dose-dependent photocleavage of NPOM from DNA as monitored *in situ* by UV-Vis absorption spectroscopy. **(A)** Absorption at 395 nm versus time. Dotted lines indicate the reaction kinetics fitted using first order exponential decay. The half-times were 54 ± 7 s for NPOM-DNA for NPOM-DNA (orange circles) and 52 ± 8 s for NPOM-dT (brown circles). **(B)** Absorption spectra of NPOM-DNA (AT2, orange), NPOM-dT (brown), and unmodified DNA duplex (AT1, blue) before (solid line) and after 120 s of light irradiation (dotted line).

### NPOM lowers the thermal stability of DNA duplex which can be reversed by photocleavage

Several studies show that Rad4/XPC-binding and NER repair propensity for various lesions are positively correlated with the thermal destabilization induced by the lesion, which can be measured by DNA melting temperatures^40, 50, 58^. To see if thermal stability of DNA was impacted by the NPOM modification, we measured the melting temperatures (T_m_) of the NPOM-DNA before and after photocleavage and compared them with that of the unmodified DNA. The T_m_ of NPOM-DNA (AT2, 45.2 °C) was ~7 °C lower than that of the unmodified DNA (AT1, 52.0 °C) while the T_m_ of NPOM-DNA after photocleavage (2 min irradiation) was the same as that of the unmodified DNA (52.0 °C) (**Table 1, Figure S2A**). These results showed that covalent NPOM adduct destabilized the DNA duplex but its photoremoval restored the DNA stability. The reaction products such as nitrosoacetophenone, though present in the reaction mix, did not affect the DNA thermal stability.

### Competitive electrophoretic mobility shift assays (EMSA) show that NPOM-DNA is specifically recognized by Rad4, which is abolished upon NPOM photocleavage

After observing NPOM is a helix-destabilizing DNA adduct, we set out to examine if the adduct can indeed be recognized as a DNA lesion by Rad4/XPC, by using a competitive electrophoretic mobility shift assay (EMSA) as extensively used before (**Figure 3**)^40–41, 43, 47, 50^. In this assay, the binding of the protein to 5 nM ^32^P-labeled substrate DNA is monitored in the presence of 1,000 nM unlabeled, undamaged “competitor” DNA (CH7_NX). The NPOM-DNA (AT2) showed ~15-fold lower apparent dissociation constant (K_d,app_~48 nM) than the corresponding unmodified DNA (AT1) (K_d,app_ ~701 nM). This specificity of NPOM-DNA (AT2) is even slightly higher than another specific model DNA substrate containing CCC/CCC mismatches (CH10_NX; K_d,app_ ~79 nM), and is comparable to that of a bona fide NER lesion, 6-4 thymidine-thymidine photoproduct (6-4PP) (K_d,app_ ~35 nM)^40^. On the other hand, the NPOM-DNA after photocleavage (AT2 + *hν*) showed K_d,app_ (~744 nM) comparable to that of the unmodified DNA (AT1). These results show that the NPOM adduct in DNA is specifically recognized by Rad4/XPC as a lesion and that its photoremoval abolishes the specific binding. Therefore, the NPOM adduct could be useful as a new photoswitchable model DNA lesion for Rad4/XPC and potentially for NER.

**Figure 3.**
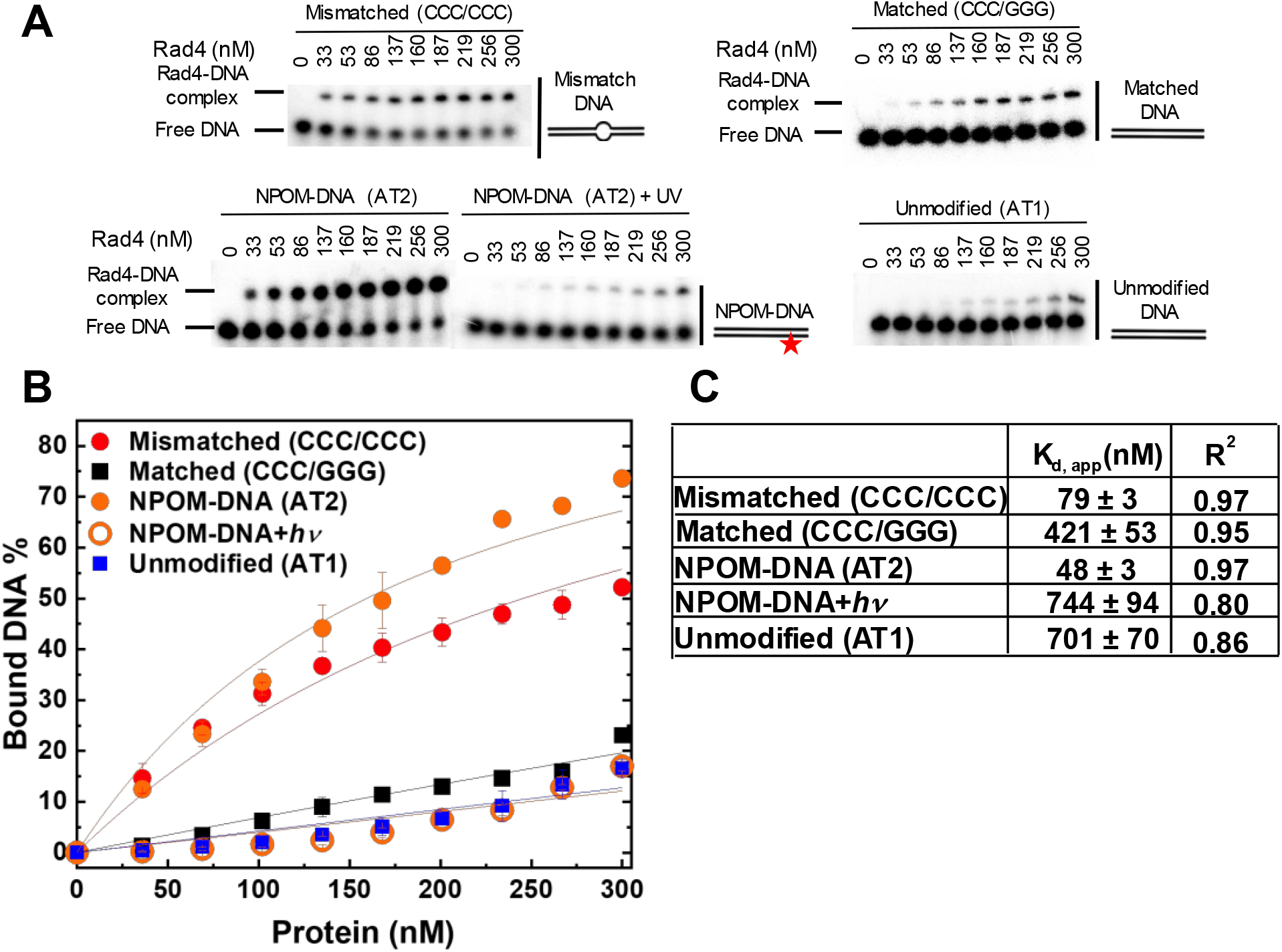
Apparent Rad4-binding affinities of DNA constructs measured by competition electrophoretic mobility shift assays (EMSA). **(A)** Typical gel images showing the wild-type Rad4-Rad23 complex binding to various DNA constructs. Mismatched (CCC/CCC) and matched (CCC/GGG) DNA represent typical specific and nonspecific binding substrates, respectively. The sequences of DNA are in Table S1. **(B)** Quantification of the Rad4-bound DNA fractions versus total concentrations of the protein from gels including those shown in (B). The symbols and error bars indicate the means and ranges as calculated by ± sample standard deviations, respectively, from triplicate experiments. Solid lines indicate the fit curves of the data point. **(C)** K_d,app_ and R^2^ of the fits derived from (B).The errors indicate the errors of the nonlinear regression fit.

### DNA conformation landscape mapped by fluorescence lifetime analyses (FLT) of the tC^o^-tC_nitro_-labeled DNA shows DNA with NPOM becomes heterogeneously distorted upon binding to Rad4

Previously, we showed that fluorescence lifetime (FLT) analyses combined with a unique set of FRET probes (tC^o^ and tC_nitro_) in DNA can be used to map the conformations of DNA in solution^43^. The tC^o^ and tC_nitro_ are a FRET pair that serve as donor and acceptor, respectively^44–45^. As cytosine analogs, these probes retain normal Watson-Crick pairing with guanines with minimal perturbation of DNA structure and stability^44, 50^. Furthermore, the rigid exocyclic ring and its base stacking interactions hold these nucleotide analogs in relatively fixed orientations within the DNA helical structure, making their FRET sensitive to subtle distortions in DNA helicity that alter the probes’ separation and/or relative orientation^59–61^. For example, Rad4-induced untwisting and ‘opening’ of 3-bp mismatched DNA could be monitored by the FRET efficiency between tC^o^ (donor) and tC_nitro_ (acceptor) placed on either side of the mismatch^43, 50^. The FRET efficiency (E) relates directly to the lifetimes of the excited donor fluorophore, as 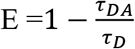, where *τ_DA_* and *τ_D_* are the donor lifetimes in the presence and absence of the acceptor, respectively. The lifetime approach offers distinct advantages over other techniques such as single-molecule FRET and is a more robust way to obtain FRET efficiency than the intensity-based steady-state measurements^43, 62^. Here, we adopted our previous approach and incorporated the tC^o^-tC_nitro_ FRET probes in the context of the NPOM-DNA construct (AT2) in the same positions relative to the lesion site as before (**Table 1**)^43, 50^. tC^o^-tC_nitro_ probes did not significantly alter DNA duplex stability of these DNA constructs as measured by DNA melting temperatures (**Figure S2B**)^43, 50^. The fluorescence decays of each sample were obtained (**Figure S3**) and analyzed using two different methods, discrete exponential (DE) and maximum entropy method (MEM), as before^43^. Results from DE analyses are shown in **Figures S4-S5** and **Table S1,** and MEM results are detailed in **Figures S5-S6** and **Tables S2**. Both analyses resulted in FLT profiles largely consistent with each other (**Figure S5**), as shown before^43^. Our discussion below is primarily based on the results obtained from MEM.

First, for the unmodified DNA (AT10), the donor-only construct (AT10_D) showed a single major lifetime peak (*τ_D_*) at 5.1 ns, which corresponds to the intrinsic lifetime of the donor fluorescence (since there is no acceptor and thus no FRET), consistent with previous results by us and the Wilhelmsson group (**Figures 4A & S5A**)^43, 63^. In comparison, the DNA containing both the donor and acceptor (AT10_DA) showed a major lifetime peak (*τ_DA_*) at 0.27 ns with 86% fractional population with minor peaks at 1.8 ns (7%) and 4.8 ns (7%) (**Figures 4A & S5B**). The major lifetime peak of ~0.3 ns corresponds to a FRET efficiency of ~0.94, which closely matches the calculated FRET of 0.936 for an ideal B-DNA structure^43, 63–64^. The 4.8 ns lifetime is close to the intrinsic lifetime of the donor in the absence of the acceptor; however, this was not due to an excess of unannealed donor strand, as the same was observed even in the presence of 50% excess acceptor strand (**Figure S7**). These characteristics of AT10_DA agree well with those of other matched DNA duplexes we had previously examined and confirm that AT10 mainly adopts B-DNA conformation with perhaps a minor population of non-B-DNA conformations^43^. Also, Rad4-binding to the DNA did not alter the FLT profile, as shown with other nonspecific DNA, indicating that nonspecific binding by Rad4 does not lead to detectable changes in tC^o^-tC_nitro_-based FRET (**Figures 4A & S5C**).

**Figure 4.**
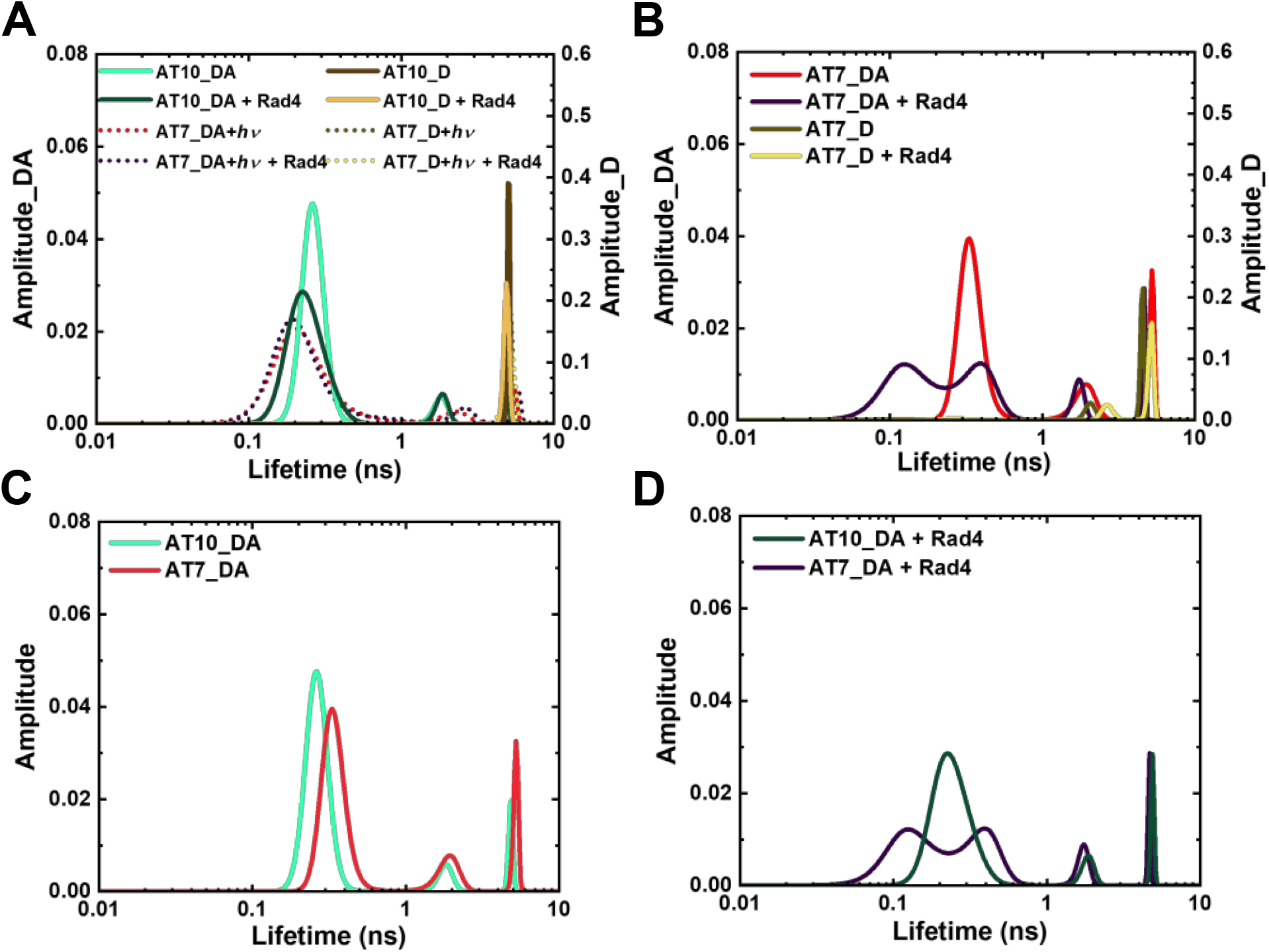
Fluorescence lifetime distributions obtained from MEM analyses for various tC^o^-tC_nitro_-labeled DNA and DNA-protein complexes. “_D” indicate DNA with donor only; “_DA” indicate DNA with donor/acceptor pair. **(A)** Unmodified DNA (**AT10**) in the absence and presence of Rad4 and its comparison with NPOM-DNA after 120 s of photocleavage reaction (**AT7+hν**). **(B)** NPOM-modified DNA (**AT7**) in the absence and presence of Rad4. **(C, D)** Overlay of unmodified (AT10_DA) and NPOM-DNA (AT7_DA) without Rad4 **(C)** and in the presence of Rad4 **(D)** Reproducibility of MEM FLT distributions for each DA sample is shown in **Figure S6**. Full reports of the lifetimes, fractional amplitudes, FRET efficiencies of each peak as well as the sample’s average FRET efficiencies are in **Table S2**.

In comparison to a single peak profile in the unmodified AT10_D, the donor-only NPOM-modified DNA (AT7_D) showed two peaks: one major peak with a lifetime of 4.5 ns, similar to the *τ_D_* of AT10, but also a minor, 2.0 ns peak (**Figures 4B & S5D**). This additional 2.0 ns peak was present even for unannealed, single-stranded AT7_D, indicating that it is not sensitive to the DNA’s conformation (**Figure S5J**), and it disappeared upon photocleavage, as seen for AT7_D irradiated for 120 s (AT7_D+*hν*), indicating an influence of NPOM on the tC^o^ fluorescence (**Figures 4A & S5G**). However, despite this minor interference by NPOM on the tC^o^ fluorescence, the donor-acceptor-labeled NPOM-DNA (AT7_DA) showed remarkable resemblance to that of the unmodified DNA (AT10_DA) with one major (0.31 ns (74%)) and two minor peaks (1.8 ns (13%) and 4.8 ns (13 %)) (**Figures 4B, 4C & S5E**). The similarity between the two DNA constructs indicates that the conformations of NPOM-modified DNA as sensed by the tC^o^-tC_nitro_ pair are largely unperturbed by the NPOM modification and most retain B-DNA-like conformation. Upon binding to Rad4, however, the FLT profile of AT7_DA changed distinctly compared with unbound DNA, unlike with AT10_DA (**Figures 4B, 4D & S5F**). Two broader and shorter lifetime peaks (0.16 ns (55%) and 0.39 ns (31%)) replaced the single major peak for unbound AT7_DA at ~0.3 ns while the 1.7 ns and 4.7 ns peaks reduced to 8% and 6% in the fractional population, compared with DNA without Rad4. Such changes in the lifetime distribution translates to an increase in the average FRET efficiency from 0.78 to 0.87 upon Rad4 binding. A broader distribution of lifetimes with multiple peaks in AT7_DA indicated that NPOM-DNA, when specifically bound to Rad4, can access a broader range of distinct conformations with some that deviate from B-DNA. However, the FRET value of the Rad4-bound DNA is significantly different from the FRET E calculated based on the ‘open’ DNA conformation as seen in the crystal structure of Rad4-bound lesions (0.043), suggesting perhaps a different binding mode for this DNA than other specific substrates^41^, which we discuss further in Discussion.

Lastly, NPOM-DNA after photocleavage (AT7+*hν*) showed profiles closely resembling that of the unmodified AT10 without or with Rad4, consistent with the expected photoconversion of NPOM-DNA to unmodified DNA (**Figure 4A & S5G-I**). The small differences in the peak positions and widths were due to the nitrosoacetophenone released after photocleavage, as such differences largely disappeared upon its removal using a G25 size-exclusion resin (**Figure S5K**). These results reaffirm that light-induced cleavage of the NPOM group from DNA abolishes the specific binding of Rad4 to the DNA and that the FLT profiles can be used to differentiate specific versus nonspecific binding of Rad4 with DNA.

#### Progressive, light-induced conversion from specific to nonspecific Rad4-DNA complexes as tracked by FLT

Seeing that FLT can discern the different conformational landscapes of NPOM-DNA when specifically bound versus nonspecifically bound to Rad4, we next examined progressive changes in FLT of Rad4-bound NPOM-DNA upon gradual increase in photoirradiation times (0-120 sec) (**Figure 2**). Progressive increase in photocleavage with increased irradiation time under these conditions resulted in little change in the FLT distributions for free, unbound NPOM-DNA: it mostly retained B-DNA conformation (*τ* = 0.3 ns) although some broadening of the peaks was observed (**Figures S8A-E**). In comparison, the FLT profiles of the Rad4-bound NPOM-DNA changed distinctly with the increased irradiation (**Figures 5A & S8F-J**). For instance, the two Gaussian peaks at ~0.16 ns and ~0.4 ns had comparable fractional amplitudes (1.9:1) before irradiation but their ratios gradually increased with irradiation (2.8:1 at 60 sec), eventually merging as a single peak with *τ* of ~0.3 ns, closely resembling the non-specifically bound unmodified AT10_DA (**Figure S8J**). Interestingly, the same tendency was observed when the ratio between specific and nonspecific binding was altered by progressive change in DNA:protein ratios (**Figures 5B & S9-S10**). The FLT profiles after 30 sec or 60 sec irradiation resemble the profiles obtained when NPOM-DNA was bound to 2- or 3-fold molar excess of Rad4 (**Figure 5C-D**). These results indicate that partial irradiation results in a mixture of specifically and nonspecifically bound complexes, as anticipated, yielding conformational distributions that are similar to when there is excess of protein and thus competition between specific and nonspecific binding.

**Figure 5.**
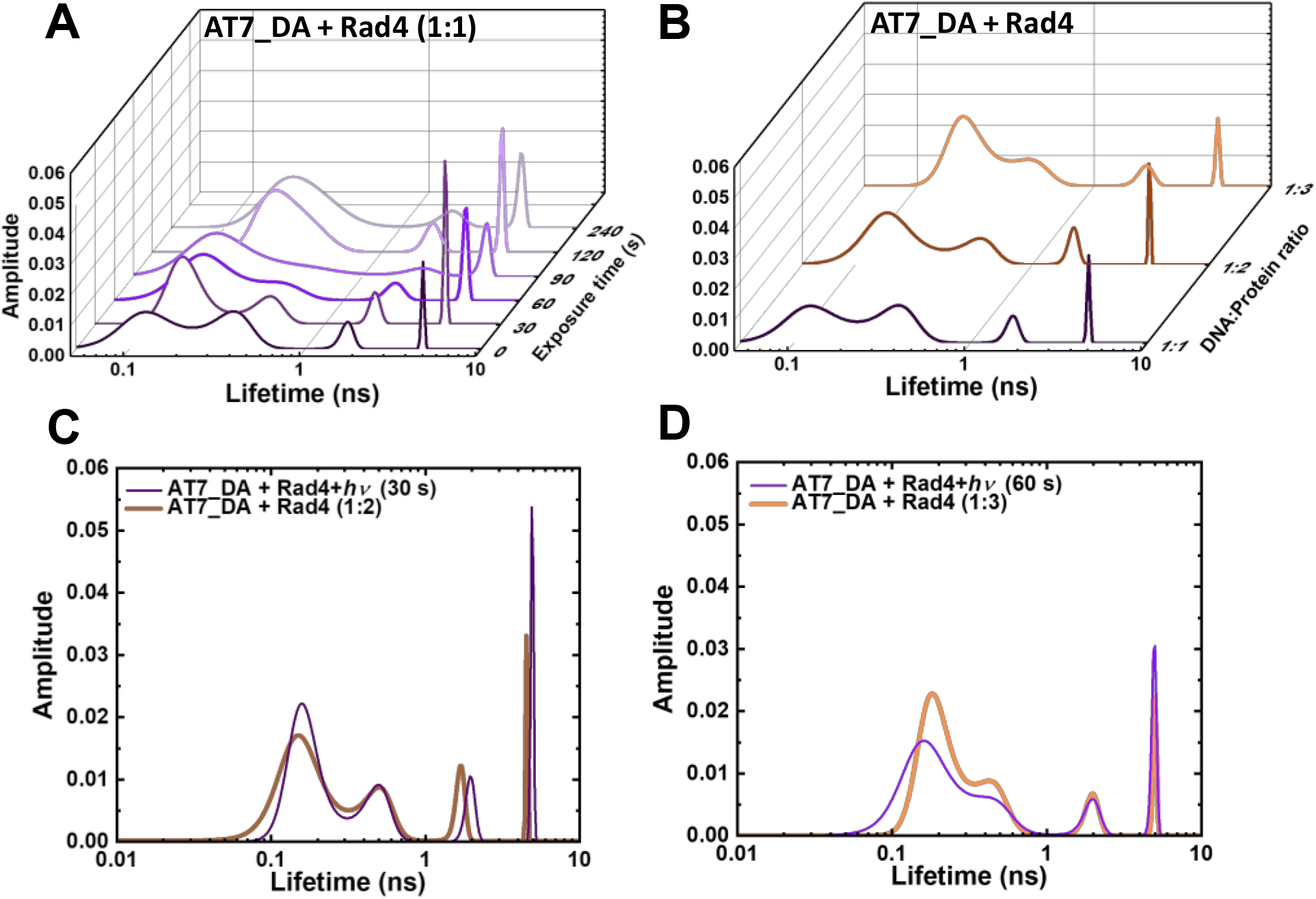
Light-induced conversion from specific to nonspecific Rad4-DNA complexes as tracked by FLT. **(A)** FLT distributions obtained from MEM analyses of Rad4-bound NPOM-DNA (AT7_DA) with varying photocleavage irradiation time (0-240 s). **(B**) FLT distributions of NPOM-DNA with varying DNA:Rad4 ratios. **(C)** Overlay of 30 s irradiation and 1:2 AT7_DA:Rad4 complex. **(D)** Overlay of 60 s irradiation and 1:3 AT7_DA:Rad4 complex.

### Conformational searches and MD simulations reveal two predominant major groove and one base-displaced intercalated conformation for the NPOM-modified DNA duplex

Our FLT study indicates that the NPOM-modified DNA retains a majority B-DNA conformation, at least as sensed by the tC^o^ and tC_nitro_ FRET probes in these constructs. To gain molecular insights into the FLT/FRET data and understand how the NPOM adduct may impact the DNA duplex structure, we turned to extensive all-atom molecular dynamics (MD) simulations on NPOM-modified DNA. As there is currently no structure available for the NPOM-adduct containing DNA, we first carried out extensive conformational searches to obtain initial models for MD simulations of NPOM-dT in B-DNA (**Scheme 1**). The search produced five geometry optimized rotamer combinations of NPOM-dT that could fit into the 13-mer B-DNA structure without causing extensive distortions to the duplex (**Figure S11**). Among these five conformations of NPOM-dT, there were four major groove conformations where NPOM adopted various orientations in the major groove with the dT in *syn* conformation (MJ1, MJ2, MJ3 and MJ4 in **Figure S12**), and one base-displaced intercalated conformation where NPOM-dT intercalated into the helix with the dT in *anti* conformation and its partner dA extruded into the major groove (INT in **Figure S12**). We carried out 1.5 μs MD simulations for each of these systems as well as an unmodified control duplex (**Figure S13**). Among our MD simulations of major groove conformations, one ensemble exhibited denaturing of the duplex and extensive distortions (MJ3 in **Figure S13A**) and hence was excluded from our further analyses of the structural ensembles.

Our stable 1.5 μs MD simulations of major groove structures (MJ1, 2 and 4) for NPOM-dT converged to two predominant conformations: two rotamers around the long axis of the NPOM rings that placed the nitro group toward the major groove surface (MJ-I) or toward the solvent (MJ-II) (**Figures 6 & S13C**). These two conformations were observed in all three stable MD ensembles with varying proportions in each population (**Figure 6A**); they were able to flexibly interchange through different combinations of rotations around the dihedral angles between the NPOM rings and the modified dT (**Figures 6 & S14**). Of the combined ensembles, 66% adopted either of these two major groove conformations. In the major groove conformation with the nitro group facing the major groove surface (MJ-I; 31% of the population), the NPOM rings were oriented along the helix axis on the major groove surface with the five-atom ring pointing toward the 5’ end of the lesion-containing strand, and its partner dA extruded moderately toward the major groove. With the nitro group facing the solvent (MJ-II; 35% of the population), the NPOM rings were oriented along the base pair planes with the five-atom ring pointing toward its partner dA, protecting the dA from solvent. The remaining 34% of the major groove population were transients that occurred during the transition between the two predominant interchanging rotamers.

**Figure 6.**
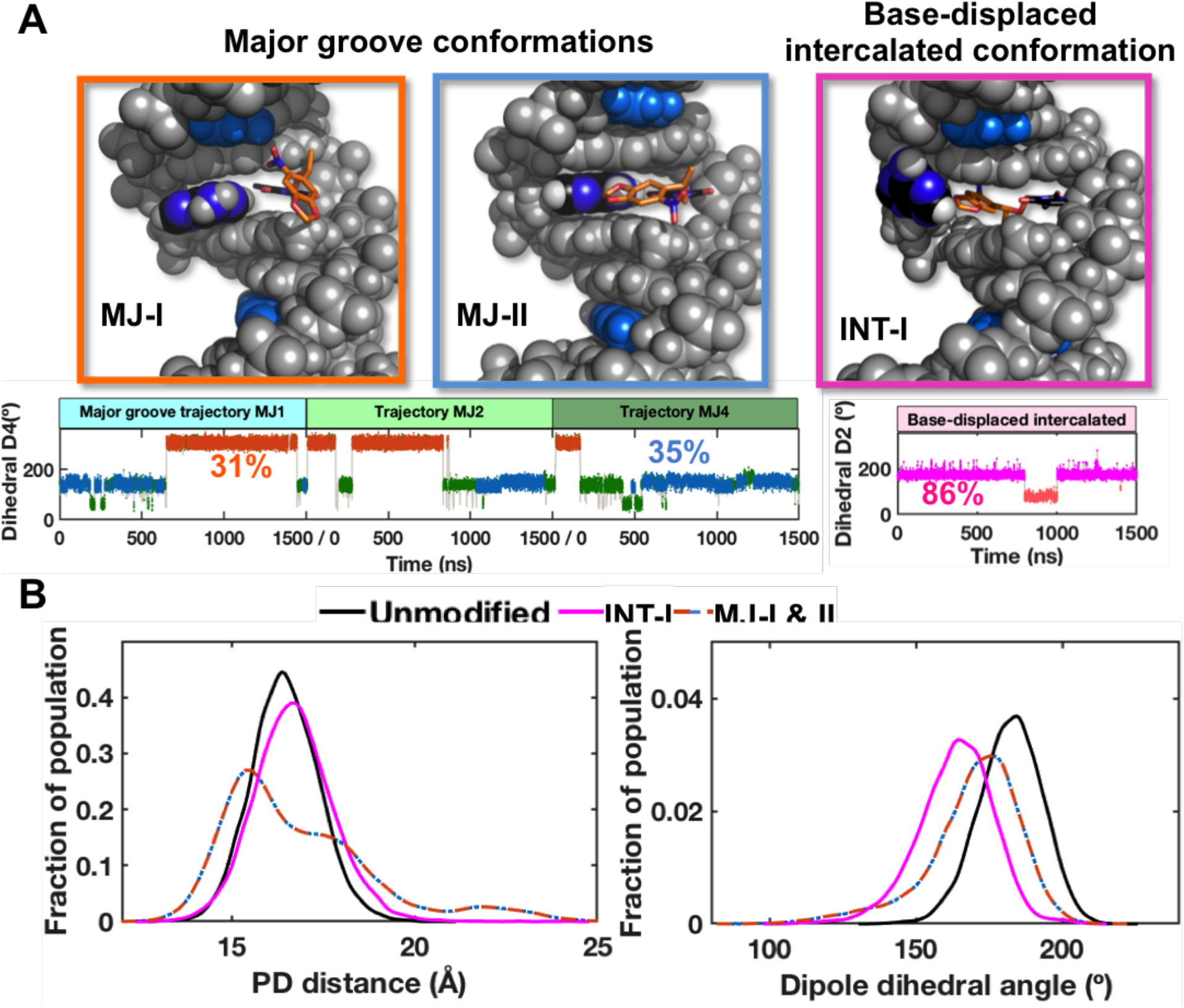
NPOM-containing DNA structures obtained from stable MD derived ensembles. **(A)** Best representative structures for the two major groove (MJ) and one base-displaced intercalated (INT) conformations. The NPOM-dT dihedral angles (**Figure S14**) that determine these different conformations are shown for each conformational family (MJ and INT) with each cluster color-coded and labeled with its percentage of population. The transient clusters in the major groove ensembles are colored green. These structures are shown in cartoon in Figure S13. **(B)** The distributions of the modeled FRET pair distances (PD distance) and dipole dihedral angles for the major groove and base-displaced intercalated NPOM-dT-containing DNA and unmodified DNA. The definitions for the PD distance and dipole dihedral angle are given in Figure S15. The major groove values are for the two dominant major groove conformations (66% of the population), and the base-displaced intercalated values are for the stable intercalated conformation (86% of the population).

For the base-displaced intercalated NPOM-dT, the structural ensemble remained stable, with the NPOM rings intercalated into the DNA helix stacked with neighboring base pairs; the partner dA was flipped into the major groove and protects the NPOM from the solvent (INT-I; **Figures 6, S13C & S14**). This intercalated conformation comprised 86% of this base-displaced intercalated conformational family. The remaining 14% represented one brief excursion during the MD simulation where the NPOM rings were folded back to stack with dT and stretched the base pair steps (INT-II; **Figure S14**).

To gain insight on the experimental FLT/FRET data, the two major groove conformations, the intercalated conformation and the unmodified duplex were further analyzed. We modeled the tC^o^-tC_nitro_ FRET pairs at the respective nucleotide positions and calculated their distances and dihedral angles between the dipoles (detailed in **SI Methods**). The distances were measured between the center of mass for the middle ring of each fluorophore model (**Figure S15A**). The dipole dihedral angles were calculated between the modeled dipoles of the FRET pair (**Figure S15B**). The distance between the FRET pairs was very similar in all conformations of the NPOM-dT-containing DNA and the unmodified DNA: 16.8 ± 1.8 Å for the major groove conformations combined for the two rotamers, 16.7 ± 0.3 Å for the intercalated one, and 16.5 ± 0.1 Å for the unmodified DNA (**Figure 6B**). While the dipole dihedral angles showed slight differences between the NPOM-dT-containing DNA and the unmodified DNA, the major groove conformations combined for the two rotamers had a value of 170 ± 11° which was close to the unmodified DNA of 182 ± 2°. However, the intercalated conformation was further from the unmodified duplex with a value of 164 ± 5° (**Figure 6B**). The FRET efficiencies based on the modeled FRET pairs were ~0.96-0.98 for the best representative structures of these conformers, not too far away from the value expected of ideal B-DNA (0.936) and consistent with the FLT experimental results.

These MD simulations provide atomistic models for the NPOM-DNA and insights into their structural dynamics. While the major groove lesion-containing DNA structures were similar to the unmodified DNA, they also exhibited local lesion conformational dynamics (**Figure 6A**) that may be relevant to the recognition of the NPOM adduct by Rad4 (see Discussion).

## DISCUSSION

Though a variety of photoswitchable DNA/RNA have been shown to modulate DNA/RNA structures and functions, their applications for elucidating DNA repair mechanisms have been relatively scarce. For instance, such chemistry has been used in DNA for inducing site-specific, single or double strand breaks^65–67^ and for triggering structural transition in a base excision repair enzyme to study its mechanism^42^. NPOM-related, 2-nitrobenzyl or 2-(2-nitrophenyl)propyl groups have also been shown to mask the recognition of mismatches by the mismatch binding protein MutS, which photo-irradiation could restore^68^. Here in this study, we show for the first time that a photoswitchable modification on DNA can be recognized as efficiently as a *bona fide* DNA lesion by a DNA damage sensor, the Rad4/XPC nucleotide excision repair complex. Furthermore, such specific binding could be abolished by photocleavage in a manner dependent on the light dose. Our FRET study also demonstrates the ability of the tC^o^-tC_nitro_ pair to detect specific binding and the loss of it upon photocleavage. This study thus sets the stage for future studies that can couple optical triggering with a variety of other techniques (e.g., fluorescence conformational dynamics spectroscopy, time-resolved x-ray crystallography, cellular repair kinetic studies, etc.).

NER is unique among DNA repair mechanisms in that it repairs an extraordinarily wide range of DNA lesions caused by various environmental and endogenous agents, including intra-strand crosslinks and bulky adduct lesions. A key to its versatility lies in its initial damage sensor protein, XPC/Rad4 that senses lesions indirectly, by detecting local thermodynamic destabilization induced by DNA damage rather than the lesion structures themselves. Our study shows that the NPOM-DNA adduct is specifically recognized by Rad4 with a specificity comparable to a bona fide NER lesion such as the 6-4 photoproduct UV lesion and that its specific binding can be abolished upon photocleavage of the NPOM-adduct. Surprisingly, our fluorescence-based conformational studies show that NPOM-adduct modification did not induce a large deviation from the canonical B-DNA form, at least as probed by the tC^o^-tC_nitro_ FRET probes positioned as in these constructs. In a previous study with DNA labeled with tC^o^-tC_nitro_ FRET probes in analogous positions, a highly specific CCC/CCC mismatch showed a broad heterogeneous distributions of fluorescence lifetimes that significantly deviated from B-DNA towards longer lifetime (lower FRET) conformations^43^. Furthermore, CCC/CCC mismatched DNA showed further decrease in the average FRET upon Rad4 binding, a direction of change in line with expected FRET changes based on the known DNA conformation in the ‘open’ crystal structure^43^. In contrast, with NPOM-DNA, we observed an increase in the average FRET upon Rad4 binding even though NPOM-DNA shows higher specific binding to Rad4 than CCC/CCC. Therefore, while both DNA substrates are specifically recognized by Rad4, our results suggest that the different specific Rad4-DNA complexes may feature significantly different DNA conformations in solution and also potentially different Rad4-binding modes for specific recognition.

Our subsequent atomistic structures obtained by MD simulation provide intriguing insights into how NPOM may be recognized by Rad4/XPC as a lesion. The existence of the two nitro group conformations in the major groove is interesting in relation to the recognition by Rad4: the electronegative nitro conformer that faces the solvent may favorably interact with positively charged Arg and Lys amino acids in the DNA binding surface of Rad4 to foster Rad4 binding to a major groove NPOM conformer. The base-displaced intercalated conformer has smaller FRET dipole dihedral angle than our unmodified control in or the major groove conformers (**Figure 6**). This smaller value represents the well-known intercalation-induced untwisting^69–70^. In comparison, untwisting is more modest in the major groove conformers. Base-displaced intercalated conformers have been shown to facilitate Rad4 recognition in computational and experimental work^71–72^. Computational studies have revealed that the displaced partner base in the case of the NER-proficient (+) *cis*-benzo[*a*]pyrene-dG adduct is readily captured by a pocket between BHD2 and BHD3 while the BHD2 hairpin binds into the minor groove and untwists the duplex; in contrast, these structural hallmarks of initial lesion recognition by Rad4 are missing when the partner nucleotide is absent, in which case the lesion becomes NER-resistant^72^. Different conformers of 2-(acetyl)aminofluorene-dG lesions have also been shown to play a role in their recognition and repair by NER^73–74^. While it is difficult to ascertain which conformation is prevalent in solution here for NPOM-DNA, base-displaced intercalated conformers can preferentially be stabilized in specific sequence contexts and be in equilibrium with major groove conformers^75–77^. Future studies are needed to reveal how Rad4-binding changes the 3D structure of NPOM-DNA and how such structures impacts its repair by NER.

## Acknowledgements

We thank the members of the Min, Broyde, and Ansari groups.

