## Supporting Information for "Light-induced modulation of DNA recognition by the Rad4/XPC damage sensor protein"

### SI Methods

#### Lesion structure modeling, optimization and initial models

The initial models of NPOM-dT were built at the center dT of a 13-mer B-DNA sequences (nucleotide steps 10 – 23, AT1 sequence **Table 1** in the main text) in either a major groove or a base-displaced intercalated conformation using Discovery Studio 2.5 (Accelrys Software Inc.). Six major groove and four base-displaced intercalated conformations that can fit in the B-DNA were obtained (**Figure S11**). The 10 candidate NPOM-dT structures were then geometry optimized in Polarizable Continuum Model (PCM) of water at HF/6-31G* level of theory using Gaussian 09[1]. Five optimized NPOM-dT structures (four in the major groove and one intercalated) were modeled into the 13-mer B-DNA without extensive distortions to the duplex, and were used as the initial models of lesion-containing DNA duplex (**Figure S12**). The 13-mer B-DNA was used as the initial model of the unmodified duplex.

#### Force field parameterization

We used ff14SB[2] and GAFF[3] force field for all MD simulations. The missing parameters for the NPOM-dT lesion were assigned utilizing similar values in GAFF force field and the values given in Myung *et al.*[4]. Electrostatic potentials for the five NPOM-dT conformations in the initial models were calculated at the HF/6-31G* level of theory using Gaussian 09[1]. The partial charges for the NPOM-dT lesion were obtained using a multiconformational restrained electrostatic potential (RESP) fit procedure[5]. The partial charges, together with atom types and additional parameters, are given in **Table S3.**

#### Water box and counterions

The initial DNA models were neutralized with $Na^+$ counterions and solvated with explicit TIP3P water[6] in a cubic periodic box with side length of 63.0 Å using the tLEAP module of the AMBER18 suite of programs [7].

#### MD simulation

All MD simulations were carried out using the AMBER18 suite of programs[7]. The Particle-Mesh Ewald method [8] with 9.0 Å cutoff for the non-bonded interactions was used in the energy minimizations and MD simulations. Minimizations were carried out in three stages. First, 500 steps of steepest descent minimization followed by 500 cycles of conjugate gradient minimization were conducted for the water molecules and counterions with a restraint force constant of 50 kcal/(mol·$Å^2$) on the solute molecules. Then, 500 steps of steepest descent minimization followed by 500 cycles of conjugate gradient minimization were carried out for the water molecules and counterions with a restraint force constant of 10 kcal/(mol·$Å^2$) on the solute molecules. In the last round, 500 steps of steepest descent minimization followed by 500 cycles of conjugate gradient minimization were carried out on the whole system without restraints. The minimized structures were then subjected to three rounds of equilibration. First, each system was equilibrated at constant temperature of 10º K for 30 ps with the solute molecules fixed with a restraint force constant of 50 kcal/(mol·$Å^2$). Then the system was heated from 10º K to 300º K over 300 ps with the solute molecules fixed with a restraint force constant of 50 kcal/(mol·$Å^2$) at constant volume. In the last round of equilibration, the restraint force constant on the solute was reduced through three steps: at 10 kcal/(mol·$Å^2$) for 200 ps, at 1 kcal/(mol·$Å^2$) for 200 ps, and then at 0.1 kcal/(mol·$Å^2$) for 200 ps with constant pressure at 300º K. Following equilibration, production MD simulations for each system were carried out in a constant-temperature, constant-pressure (NPT) ensemble at 300º K and constant pressure of 1 Atm for 1.5 μs. The temperature was controlled with a Langevin thermostat [9] with a 5 $ps^{-1}$ collision frequency. The pressure was maintained with the Berendsen coupling method [10]. A 2.0 fs time step and the SHAKE algorithm [11] were applied in all MD simulations. A 1 kcal/mol restraint was applied to the end base pair hydrogen bond donor and acceptor atom pairs during the production MDs.

**Structural analyses**

*Lesion containing 6-mer RMSD.* In order to evaluate the overall stability of the 6-mer sequence that includes the base pair steps for the FRET pairs at two ends (nucleotide steps 14 – 19, AT1 sequence **Table 1** in the main text), we calculated the RMSD for the heavy atoms of the 6-mer sequence excluding the lesion-containing nucleotide. The structural ensembles were fitted to the first frame of each MD trajectory at the heavy atoms of the two nucleotide steps on each end of the 6-mer sequence. The 6-mer RMSD were calculated for the heavy atoms of the 6-mer excluding the lesion-containing nucleotide without fitting using cpptraj module of AMBER18[7]. During the MD simulations, three trajectories started from varying major groove conformations and one trajectory started from a base-displaced intercalated conformation achieved stable conformation for the 6-mer sequence. One trajectory started from a major groove conformation exhibited ruptured lesion site and extensive distortions to the 6-mer (**Figure S13**), and hence was excluded from further structural analyses.

*Lesion conformation clustering.* We measured 4 torsion angles that determine the lesion conformation (**Figure S14**) using cpptraj module of AMBER18[7]. The D1, D2, D3, and D4 torsion angles were clustered into 2, 3, 3, and 3 groups respectively using k-means method with Euclidean distance. 42 unique combinations of all torsion angle clusters were obtained. The top 10 clusters included over 96% of total population and each cluster had a population fraction of at least 1% (**Figure S14**). Two major conformational clusters repeatedly showed up in all trajectories for the major groove conformations and were collected for further analyses.

*Best representative structures.* The best representative structure for each clustered conformational ensemble is defined as the one frame that has the lowest 6-mer RMSD value (see *Lesion containing 6-mer RMSD*) to all other frames.

*FRET efficiency calculation.* In order to estimate the distance and angle between designed FRET pair and the efficiency values, we modeled the rings of the FRET pair ($tC^o$ and $tC_{nitro}$) into the DNA duplexes. We used a $tC^o$ structure (PDB ID: 3QNO[12]) to mimic the $tC_{nitro}$ rings and superposed it to the cytosine ring at the acceptor position. We replaced the dG at the donor position with a dC by superposition at the sugar ring and then superposed another $tC^o$ to the cytosine ring at this position. The distance between the FRET pair ("PD distance") is defined as the distance between the center of mass (COM) for the middling ring of each $tC^o$ model. The dihedral angles between the dipoles of the FRET pair ("dipole dihedral angle") were obtained by adjusting the dihedral angle between the y axes of the two base pair planes with relative angles between each dipole and its y axis, and reflects the twist between the base pair steps. The vectors used for the calculation of FRET efficiency were defined as in Preus *et al.*[13]

*Block average analyses.* The PD distance and dipole dihedral angle for the two prominent major-groove and one base-displaced intercalated NPOM-dT ensembles and the unmodified DNA ensemble were analyzed using the block averaging method [14-15]. In brief, the time series data were divided into "blocks" with a block size that exceeds the longest correlation time, 20 ns in these cases. The average for each block was computed and termed "block average". The mean values and the standard deviations of the block averages were used to represent the average and the variance of averages.

Molecular structures were rendered using PyMOL 1.3.x (Schrodinger, LLC.). All MD simulation data were plotted using MATLAB 7.10.0 (The MathWorks, Inc.).

### SI Tables S1-S2

**Table S1.** Fluorescence lifetimes and FRET efficiencies from DE analyses.

| | $\tau_1$ | $A_1$ | $E_1$ | $\tau_2$ | $A_2$ | $E_2$ | $\tau_3$ | $A_3$ | $E_3$ | $\tau_4$ | $A_4$ | $E_4$ | <E> |
|---|---|---|---|---|---|---|---|---|---|---|---|---|---|
| **AT7_DA** | 0.280 ± 0.002 | 0.790 ± 0.018 | 0.944 ± 0.000 | | | | 1.410 ± 0.107 | 0.119 ± 0.009 | 0.722 ± 0.021 | 4.873 ± 0.081 | 0.089 ± 0.009 | 0.042 ± 0.016 | 0.837 ± 0.012 |
| **AT7_DA + Rad4** | 0.166 ± 0.067 | 0.642 ± 0.046 | 0.967 ± 0.013 | 0.528 ± 0.103 | 0.234 ± 0.066 | 0.896 ± 0.020 | 2.299 ± 0.106 | 0.080 ± 0.010 | 0.548 ± 0.020 | 5.260 ± 0.039 | 0.043 ± 0.011 | -0.033 ± 0.007 | 0.873 ± 0.025 |
| **AT7_DA + *hν*** | 0.266 ± 0.006 | 0.850 ± 0.000 | 0.946 ± 0.001 | | | | 1.871 ± 0.152 | 0.083 ± 0.005 | 0.632 ± 0.030 | 5.333 ± 0.137 | 0.066 ± 0.005 | -0.048 ± 0.029 | 0.854 ± 0.001 |
| **AT7_DA + *hν* + Rad4** | 0.267 ± 0.004 | 0.856 ± 0.005 | 0.947 ± 0.008 | | | | 2.057 ± 0.070 | 0.086 ± 0.005 | 0.595 ± 0.013 | 5.657 ± 0.140 | 0.056 ± 0.005 | -0.111 ± 0.027 | 0.857 ± 0.002 |
| **AT10_DA** | 0.251 ± 0.008 | 0.867 ± 0.005 | 0.950 ± 0.001 | | | | 1.930 ± 0.052 | 0.073 ± 0.005 | 0.620 ± 0.010 | 5.139 ± 0.049 | 0.059 ± 0.000 | -0.009 ± 0.009 | 0.869 ± 0.004 |
| **AT10_DA + Rad4** | 0.261 ± 0.005 | 0.847 ± 0.004 | 0.948 ± 0.001 | | | | 2.004 ± 0.072 | 0.079 ± 0.005 | 0.606 ± 0.014 | 5.216 ± 0.030 | 0.073 ± 0.005 | -0.024 ± 0.006 | 0.850 ± 0.006 |

| DNA_D | <$\tau_D$> |
|---|---|
| **AT7_D** | 3.847 ± 0.132 |
| **AT7_D + *hν*** | 5.129 ± 0.049 |
| **AT10_D** | 5.083 ± 0.017 |

The errors indicate the s.d. of the data points from three independent samples.

**Table S2.** Fluorescence lifetimes and FRET efficiencies from MEM analyses

| | $\tau_1$ | $A_1$ | $E_1$ | $\tau_2$ | $A_2$ | $E_2$ | $\tau_3$ | $A_3$ | $E_3$ | $\tau_4$ | $A_4$ | $E_4$ | <E> |
|---|---|---|---|---|---|---|---|---|---|---|---|---|---|
| **AT7_DA** | **0.309 ± 0.006** | **0.742 ± 0.037** | **0.939 ± 0.001** | | | | **1.768 ± 0.087** | **0.125 ± 0.012** | **0.653 ± 0.017** | **4.806 ± 0.075** | **0.131± 0.026** | **0.058 ± 0.014** | **0.785 ± 0.028** |
| **AT7_DA + Rad4** | **0.157 ± 0.015** | **0.545 ± 0.029** | **0.969 ± 0.003** | **0.391 ± 0.007** | **0.306 ± 0.035** | **0.923 ± 0.001** | **1.705 ± 0.025** | **0.082 ± 0.002** | **0.665 ± 0.005** | **4.673 ± 0.006** | **0.065 ± 0.005** | **0.080 ± 0.006** | **0.870 ± 0.004** |
| **AT7_DA + $h\nu$** | **0.214 ± 0.014** | **0.892 ± 0.007** | **0.957 ± 0.002** | | | | **2.237 ± 0.051** | **0.052 ± 0.008** | **0.561 ± 0.010** | **5.363 ± 0.064** | **0.054 ± 0.000** | **-0.051 ± 0.012** | **0.874 ± 0.004** |
| **AT7_DA + $h\nu$ + Rad4** | **0.202 ± 0.009** | **0.902 ± 0.003** | **0.960 ± 0.001** | | | | **2.414 ± 0.145** | **0.053 ± 0.007** | **0.526 ± 0.028** | **5.502 ± 0.308** | **0.043 ± 0.007** | **-0.078 ± 0.060** | **0.877 ± 0.002** |
| **AT10_DA** | **0.265 ± 0.000** | **0.863 ± 0.000** | **0.948 ± 0.000** | | | | **1.830 ± 0.007** | **0.068 ± 0.000** | **0.641 ± 0.001** | **4.846 ± 0.005** | **0.068 ± 0.000** | **0.050 ± 0.001** | **0.863 ± 0.000** |
| **AT10_DA + Rad4** | **0.231 ± 0.011** | **0.855 ± 0.009** | **0.954 ± 0.002** | | | | **1.841 ± 0.097** | **0.065 ± 0.003** | **0.639 ± 0.019** | **4.915 ± 0.048** | **0.076 ± 0.002** | **0.036 ± 0.009** | **0.861 ± 0.002** |

| **DNA_D** | **$\langle\tau_D\rangle$** |
|---|---|
| **AT7_D** | **3.773 ± 0.066** |
| **AT7_D + $h\nu$** | **5.142 ± 0.012** |
| **AT10_D** | **5.102 ± 0.078** |

The errors indicate the s.d. of the data points from three independent samples.

**Table S3.** Atom names, types, topologies, partial charges and added force field parameters for the NPOM-dT lesion.

| Name | Tpye | Topology | Partial charge |
|---|---|---|---|
| P | P | M | 1.440501 |
| OP1 | O2 | E | -0.787291 |
| OP2 | O2 | E | -0.787291 |
| O5' | OS | M | -0.625530 |
| C5' | CT | M | 0.067372 |
| H5'1 | H1 | E | 0.054210 |
| H5'2 | H1 | E | 0.054210 |
| C4' | CT | M | 0.108084 |
| H4' | H1 | E | 0.098912 |
| O4' | OS | S | -0.341532 |
| C1' | CT | B | 0.162803 |
| H1' | H2 | E | 0.082150 |
| N1 | N* | B | -0.031648 |
| C2 | C | S | 0.428169 |
| O2 | O | E | -0.454600 |
| C6 | CM | B | -0.266057 |
| H6 | H4 | E | 0.227272 |
| C5 | CM | B | 0.009607 |
| C7 | CT | 3 | -0.248599 |
| H71 | HC | E | 0.084802 |
| H72 | HC | E | 0.084802 |
| H73 | HC | E | 0.084802 |
| C4 | C | B | 0.499678 |
| O4 | O | E | -0.509627 |
| N3 | N* | S | -0.113688 |
| CN7 | CT | 3 | 0.047549 |
| HN10 | HC | E | 0.093631 |
| HN11 | HC | E | 0.093631 |
| ON3 | OS | S | -0.325625 |
| CN8 | CT | 3 | 0.241184 |
| HN9 | HC | E | 0.045944 |
| CN16 | CT | 3 | -0.237315 |
| HN1 | HC | E | 0.077286 |
| HN2 | HC | E | 0.077286 |
| HN3 | HC | E | 0.077286 |
| CN9 | CA | S | -0.019163 |
| CN14 | CA | B | -0.376606 |
| HN7 | HA | E | 0.207499 |
| CN13 | CA | S | 0.372688 |
| ON5 | OS | S | -0.383059 |
| CN15 | CT | 3 | 0.244021 |
| HN5 | HC | E | 0.095519 |
| HN6 | HC | E | 0.095519 |
| ON4 | OS | S | -0.378980 |
| CN12 | CA | S | 0.300725 |
| CN11 | CA | B | -0.408040 |
| HN8 | HA | E | 0.260708 |
| CN10 | CA | S | 0.004866 |
| NX | no | B | 0.778735 |
| OX1 | o | E | -0.475488 |
| OX2 | o | E | -0.423428 |
| C3' | CT | M | 0.176438 |
| H3' | H1 | E | 0.073690 |
| C2' | CT | B | -0.131850 |
| H2'1 | HC | E | 0.072944 |
| H2'2 | HC | E | 0.072944 |
| O3' | OS | M | -0.672050 |

---------------------------------

Added force field parameters

BOND

| | $K_r$ | $r_{eq}$ |
|---|---|---|
| CA-no | 322.6 | 1.468 |
| CA-OS | 372.4 | 1.373 |

ANGL

| | $K_r$ | $\theta_{eq}$ |
|---|---|---|
| CA-no-o | 68.7 | 118.10 |
| CA-CA-no | 66.9 | 119.54 |
| CA-CA-OS | 69.8 | 119.20 |
| HC-CT-OS | 50.9 | 108.70 |
| CT-OS-CA | 62.4 | 117.60 |
| CA-CT-OS | 67.7 | 110.51 |
| HC-CT-N* | 49.9 | 109.50 |
| CM-C -N* | 70.0 | 114.10 |
| C -N*-C | 70.0 | 126.40 |
| N*-C -N* | 70.0 | 115.40 |

DIHE

| | # of path | $V_n/2$ | $\gamma$ | n |
|---|---|---|---|---|
| CA-CA-no-o | 4 | 3.68 | 180.0 | 2.000 |
| X -CA-OS-X | 2 | 1.80 | 180.0 | 2.000 |

IMPR

| | $V_n/2$ | $\gamma$ | n |
|---|---|---|---|
| CA-o -no-o | 7.28 | 180.0 | 2.000 |

### SI Figures S1-S15

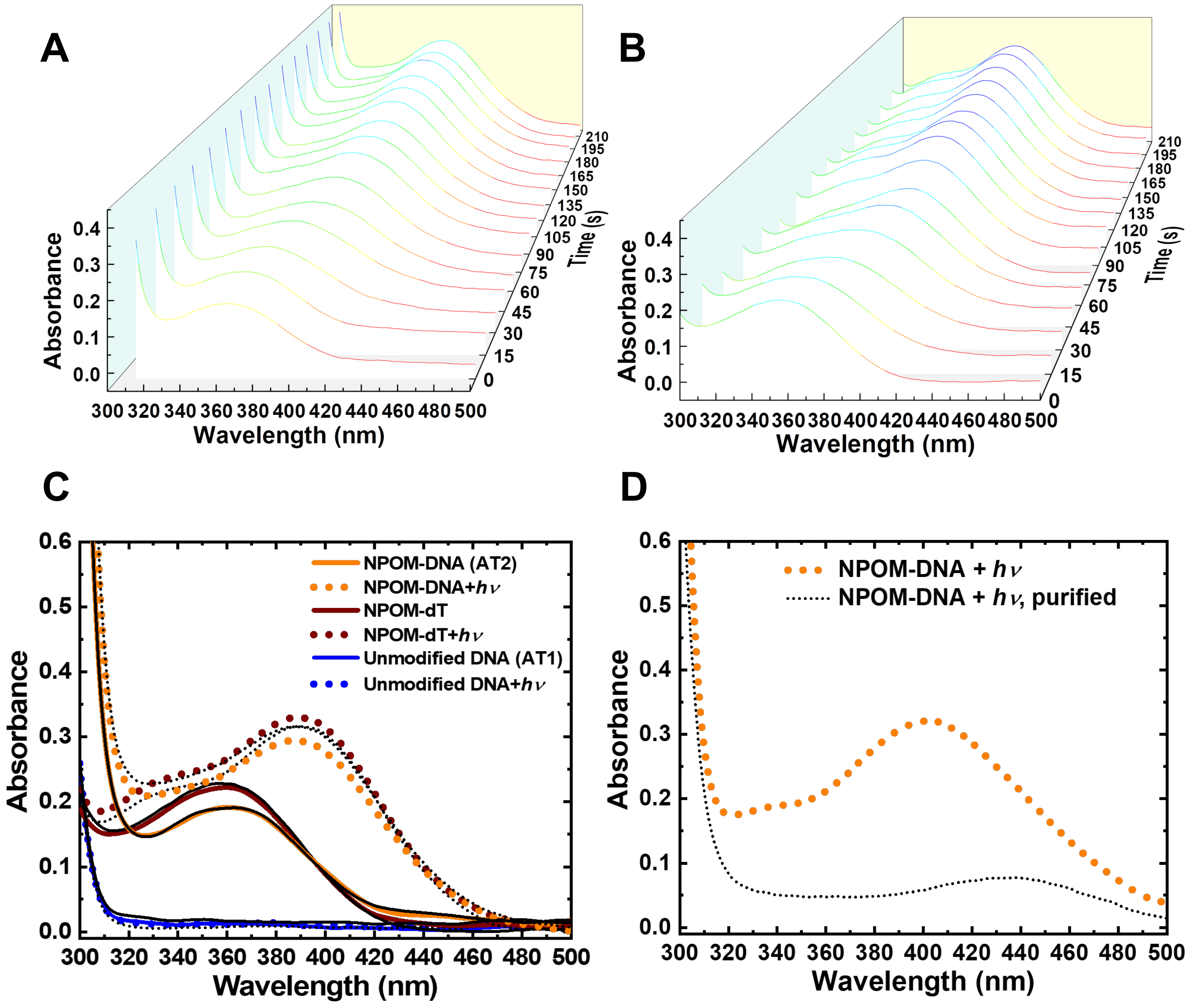


**Figure S1**. **UV/visible absorption spectroscopy for in situ monitoring of photocleavage of NPOM from DNA and dT nucleoside (A-B)** Absorption spectra of NPOM-modified DNA duplex (NPOM-DNA or AT2) **(A)** and NPOM-dT **(B)** were recorded in intervals of 15 s for 210 s and are shown in 3D graphs. **(C)** Reproducibility of the UV-Vis spectra of NPOM-DNA and NPOM-dT before and after photocleavage. Absorption spectra of NPOM-DNA (AT2, orange), NPOM-dT (brown), and unmodified DNA duplex (AT1, blue) before (solid line) and after 120 s of light irradiation (dotted line). Black indicates data from duplicate experiments. **(D)** Absorption spectra of NPOM-DNA after UV and purified with G-25 size exclusion resin.

**A**

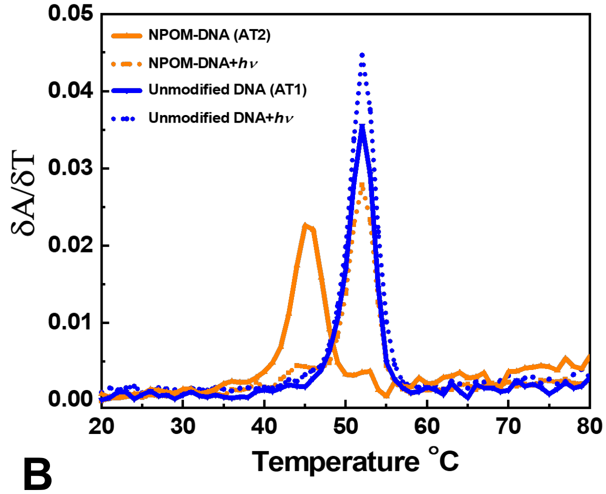


| DNA | $T_m$ (°C) |
|---|---|
| NPOM-DNA (AT2) | 45.2 ± 0.2 |
| NPOM-DNA+*hν* | 52.0 ± 0.0 |
| Unmodified DNA (AT1) | 52.0 ± 0.0 |
| Unmodified DNA+*hν* | 52.0 ± 0.1 |

**B**

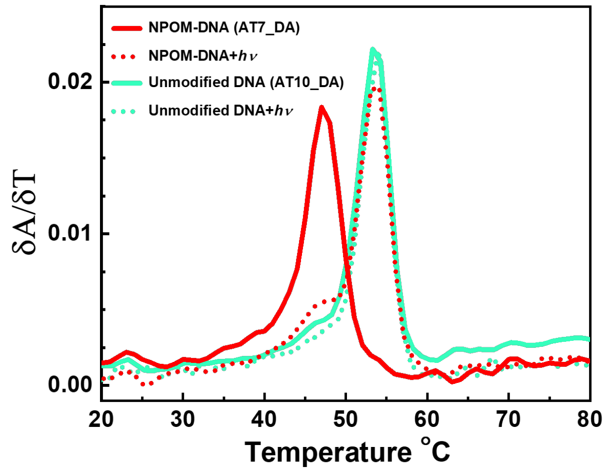


| DNA | $T_m$ (°C) |
|---|---|
| NPOM-DNA* (AT7_DA) | 47.1 ± 0.1 |
| NPOM-DNA*+*hν* | 54.0 ± 0.0 |
| Unmodified DNA* (AT10_DA) | 53.6 ± 0.5 |
| Unmodified DNA*+*hν* | 54.0 ± 0.0 |

**Figure S2**. **Thermal melting profiles of DNA duplexes before and after photocleavage.** (*Left*) The first derivatives of the absorbance at 260 nm with respect to temperature (δA/δT) versus temperature. The temperatures at the peak were designated as the $T_m$. (*Right*) $T_m$ for each construct. **(A)** NPOM-DNA (**AT2**, orange) and the unmodified DNA (**AT1**, blue) before and after irradiation (+hν) in solid and dashed lines, respectively. **(B)** DNA with tC°-$tC_{nitro}$ FRET probes: NPOM-DNA* (**AT7**, red) and the unmodified DNA* (**AT10**, green). DNA sequences for each construct are in Table 1.

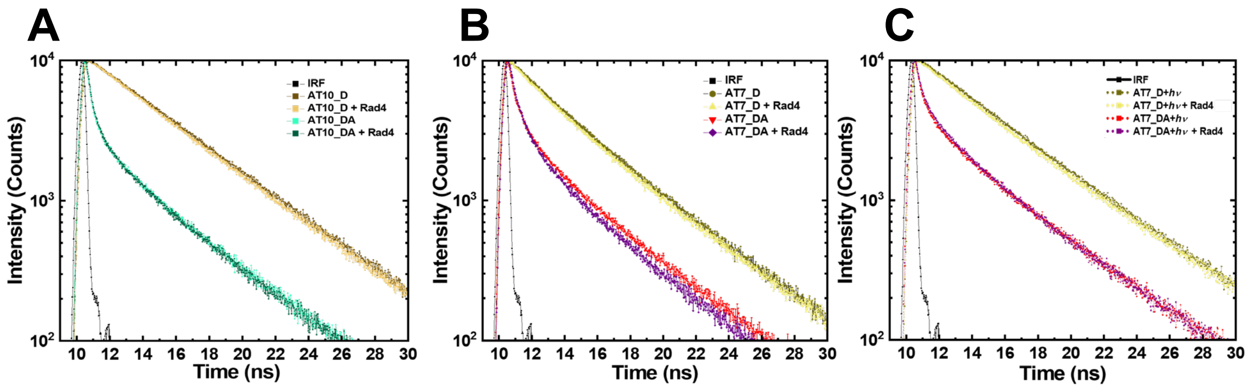


**Figure S3. Fluorescence intensity decay curves for DNA_DA and DNA_D with excitation of donor** (tC$^{o}$)**.** Donor/acceptor-labeled DNA_DA without/with Rad4 are shown for **(A)** unmodified DNA (AT10; green/dark green), **(B)** NPOM-DNA (AT7; red/purple) and (**C**) NPOM-DNA after photocleavage (AT7+hv; dotted red/dotted purple). The decay curves for corresponding donor-only labeled DNA_D are in olive/yellow. The instrument response function (IRF) is shown in black.

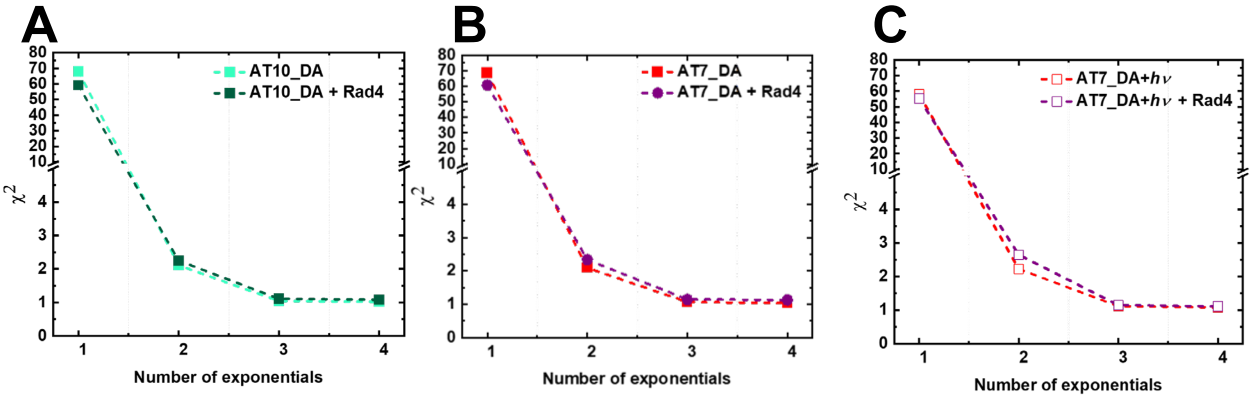


**Figure S4. The sum of the residuals $\chi^2$ as a function of increasing number of discrete exponentials.** The χ2 values (defined as $\frac{1}{N}\sum_i \frac{(y_i - y_{fit,i})^2}{\sigma_i^2}$) from a discrete exponential fit to the fluorescence decay profiles versus the number of discrete exponentials used are shown for DNA_DA samples in **(A)** unmodified DNA (AT10_DA) without and with Rad4 (green/dark green), **(B)** NPOM-DNA (AT7_DA) without and with Rad4 (red/purple) and **(C)** NPOM-DNA after photocleavage (AT7_DA + $h\nu$) without and with Rad4 (dotted red/dotted purple).

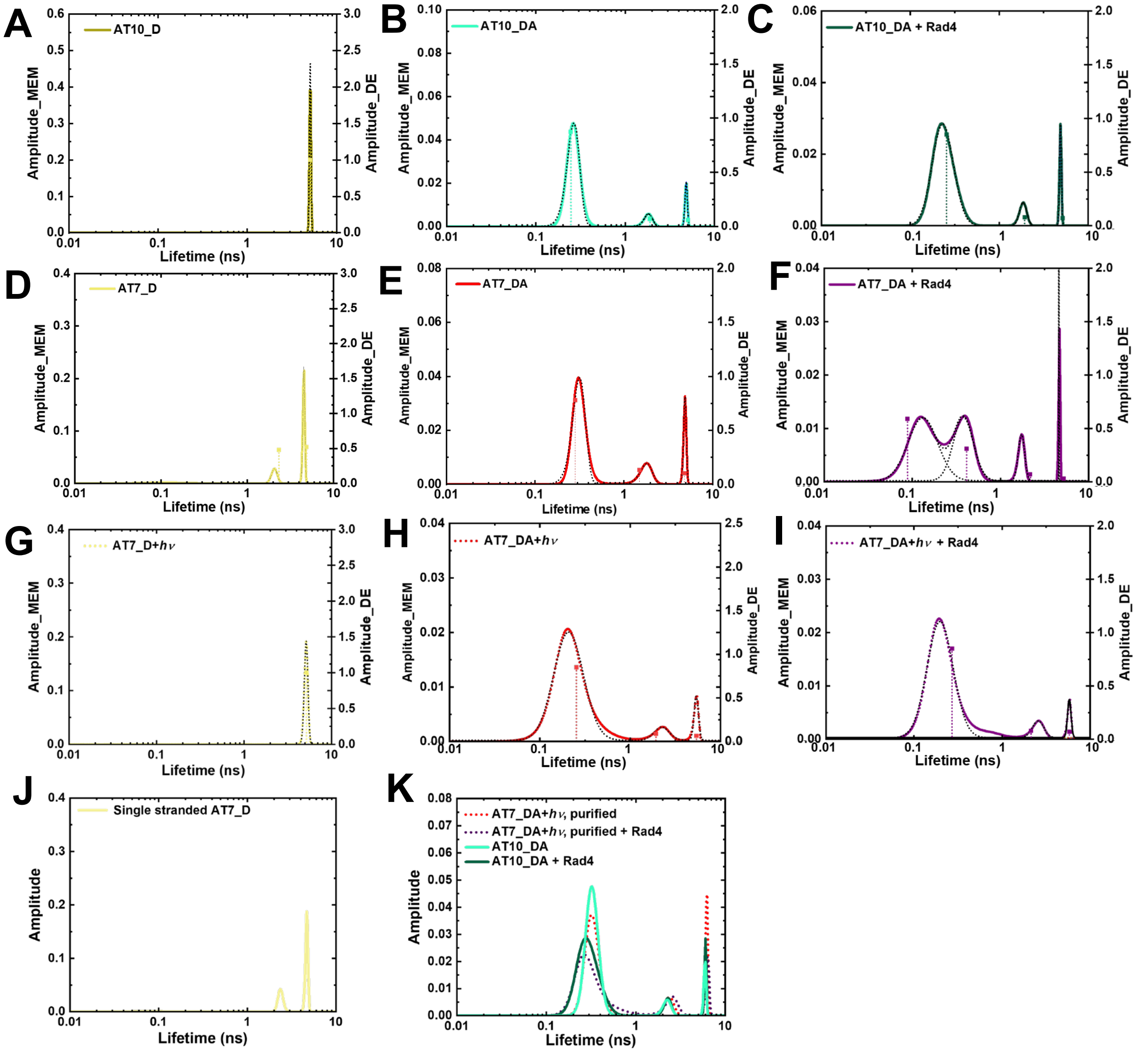


**Figure S5. Fluorescence lifetime distributions obtained from maximum entropy method (MEM)** are shown together with the lifetimes from discrete exponential (DE) analysis (vertical lines). The fractional amplitudes from the MEM (DE) are indicated on the left (right) y-axis. The Gaussian fittings for MEM distributions are shown in black dotted lines. **(A-C)** Unmodified DNA (AT10), **(D-F)** NPOM-DNA (AT7) and **(G-I)** NPOM-DNA after photocleavage (AT7+$h\nu$). (*A,D,G*) donor-only DNA (DNA_D); (*B,E,H*) donor/acceptor-labeled DNA (DNA_DA) without Rad4; (*C,F,I*) donor/acceptor-labeled DNA with equimolar Rad4. **(J)** single stranded AT7_D; **(K)** AT7_DA+$h\nu$ purified with G25 size exclusion resin, unbound and bound to Rad4. Data for AT10_DA without and with Rad4 are shown for comparison.

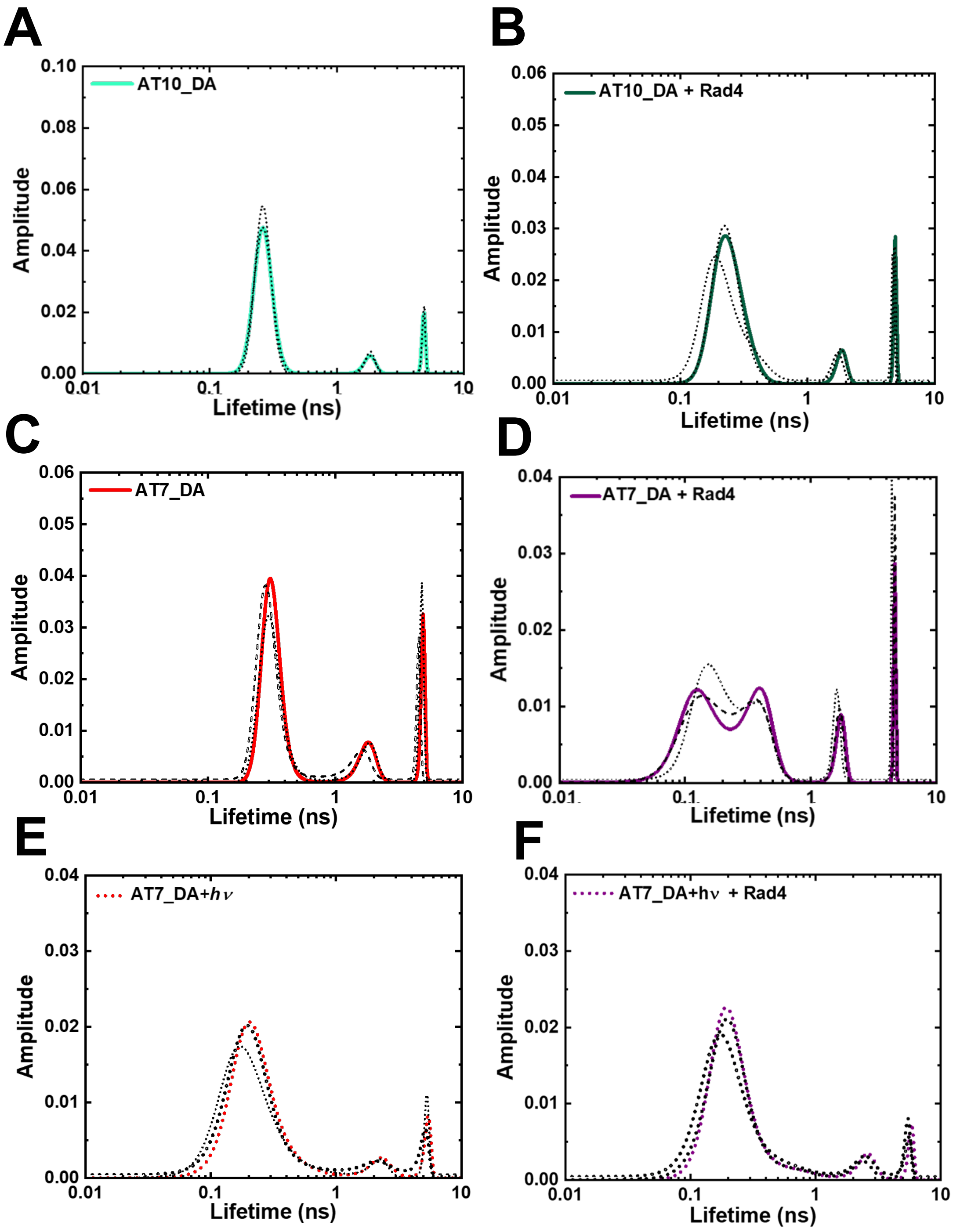


**Figure S6. Reproducibility of fluorescence lifetime distributions obtained from MEM analyses**. The MEM lifetime distributions from three independent measurements are overlaid in each panel for **(A, B)** unmodified DNA (AT10_DA) in the absence and presence of Rad4, **(C, D)** NPOM-DNA (AT7_DA) in the absence and presence of Rad4, and **(E, F)** NPOM-DNA after photocleavage (AT7_DA + $h\nu$) in the absence and presence of Rad4.

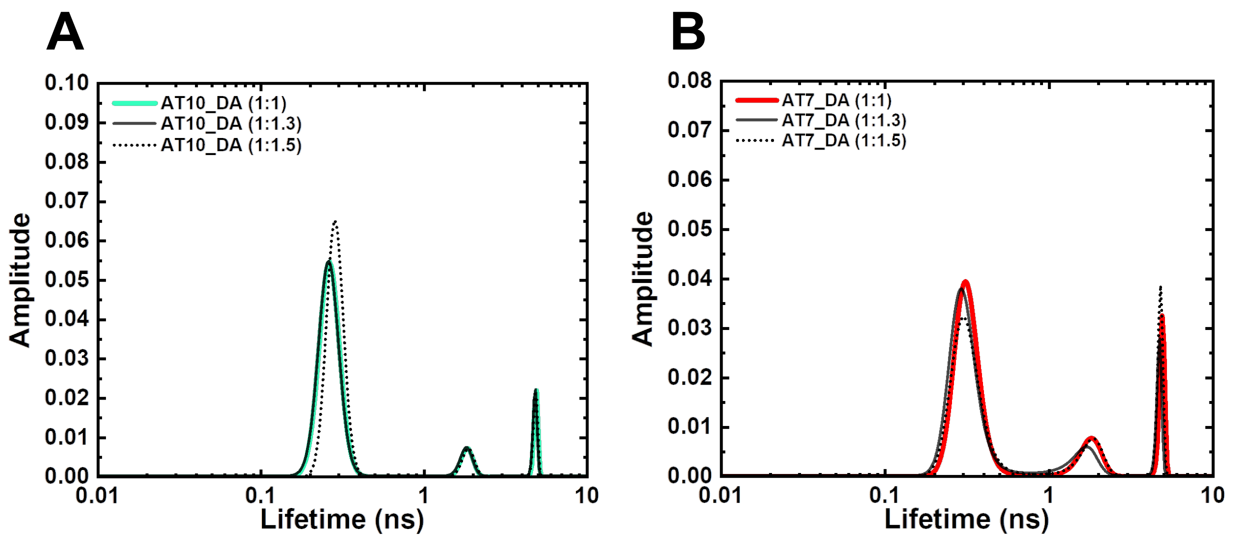


**Figure S7. FLT distributions for donor/acceptor-labeled DNA annealed with varying donor:acceptor strand ratios** for **(A)** unmodified DNA (AT10_DA, green), **(B)** NPOM-DNA (AT7_DA, red). Donor and acceptor strands are the bottom and top strands of the DNA sequences shown in Table 1, respectively. The FLT distributions remain the same among these samples indicating that the low/zero FRET peak is not due to an excess, unannealed donor strand in the samples.

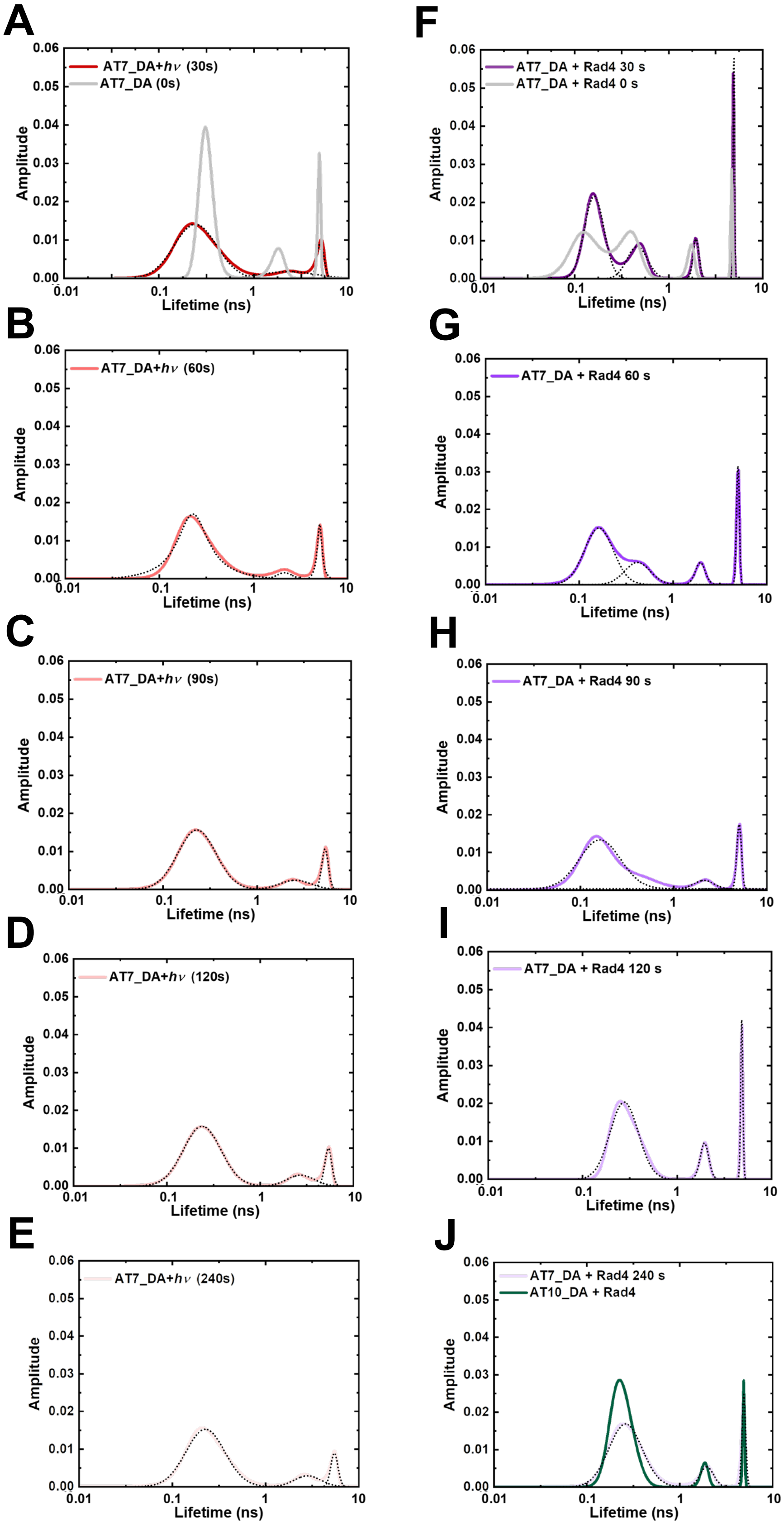


**Figure S8. FLT-MEM distributions of NPOM-DNA with varying photocleavage irradiation times**. NPOM-DNA alone (**A-E**; AT7_DA) or in the presence of equimolar Rad4 (**F-J**; AT7_DA+Rad4). **(A, F)** 0 s and 30 s **(B, G)** 60 s, **(C, H)** 90 s, **(D, I)** 120s, and **(E, J)** 240 s of photo-irradiation**.** (J) also shows unmodified DNA (AT10_DA) + Rad4 in green for comparison.

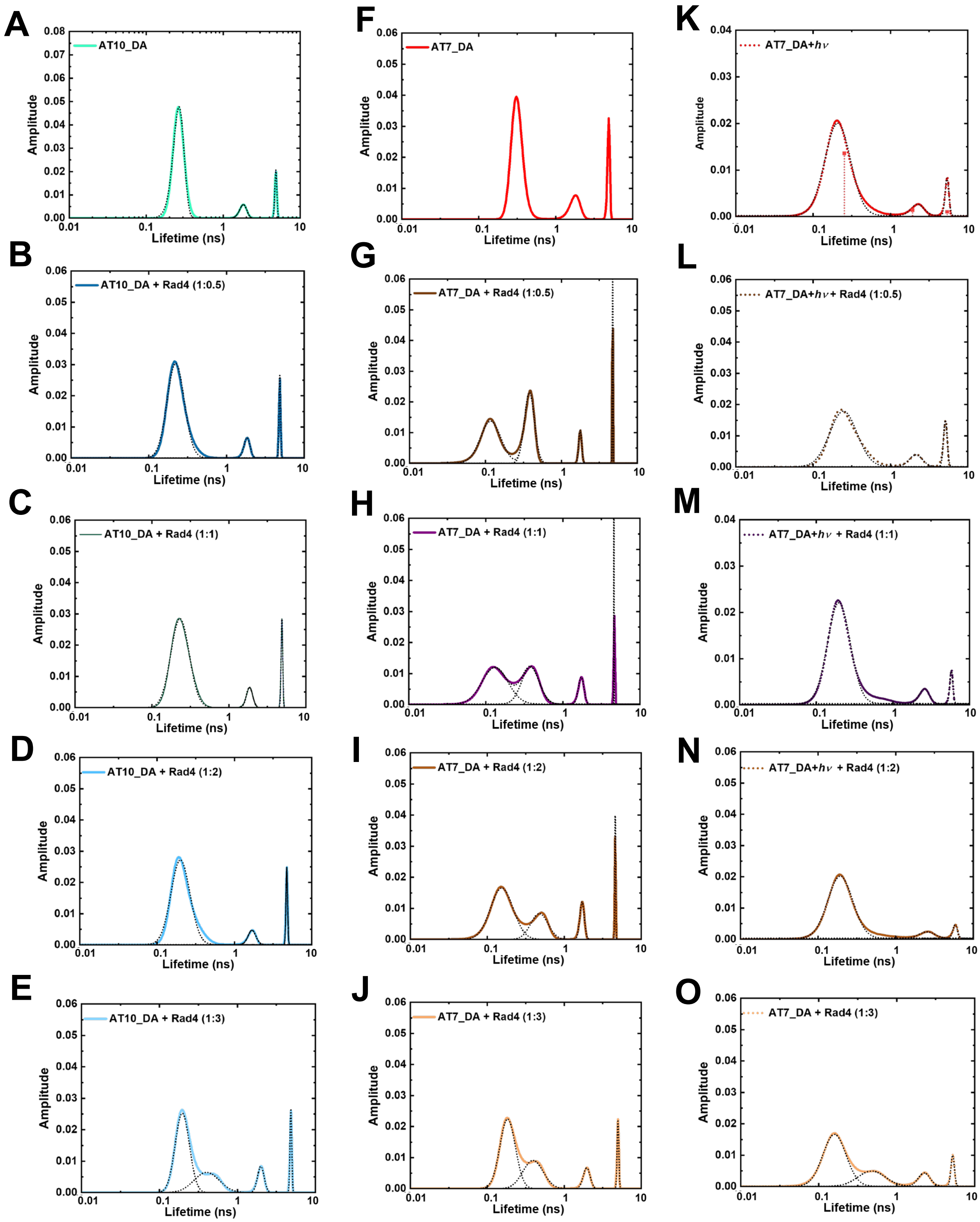


**Figure S9. MEM lifetime distributions for samples prepared with varying DNA:Rad4 ratios**. **(A-E)** Unmodified DNA (AT10_DA):Rad4 ratios were varied as **(A)** 1:0, **(B)** 1:0.5 **(C)** 1:1, **(D)** 1:2, and **(E)** 1:3. **(F-J)** show corresponding data for NPOM-DNA (AT7_DA) and **(K-O)** for NPOM-DNA after photocleavage (AT7_DA+*hν*).

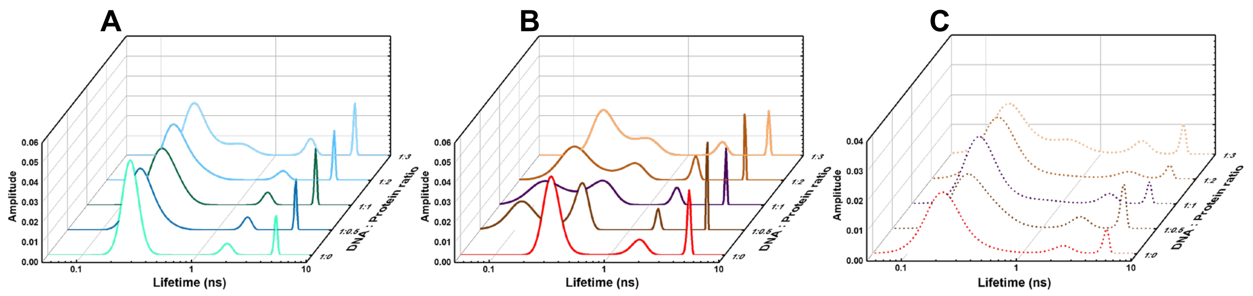


**Figure S10. MEM lifetime distributions for samples prepared with varying DNA:Rad4 ratios.** Data shown in Figure S11 are presented in 3-D graphs.

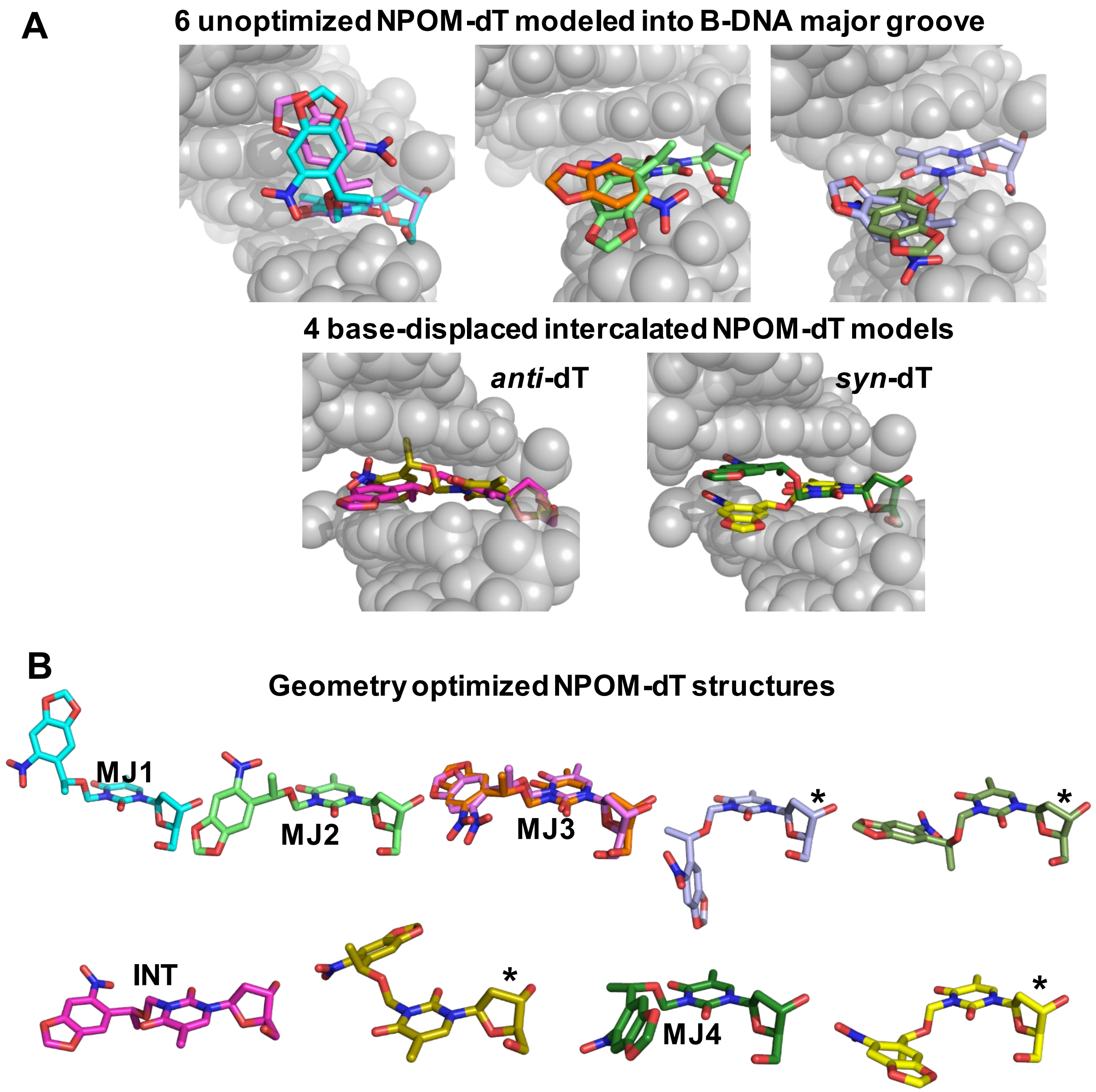


**Figure S11. NPOM-dT nucleoside conformational searches and geometry optimized conformations**. **(A)** Unoptimized NPOM-dT modeled in duplex B-DNA (gray spheres) as resulted from Stage 1 (see **Scheme 1**). NPOM-dT is shown in sticks with carbon atoms color-coded differently for each model. **(B)** Geometry-optimized NPOM-dT structures as resulted from Stage 2. Upon geometry optimization, two models converged (orange and light pink, MJ3). The NPOM-dT structures (MJ1, MJ2, MJ3, MJ4 and INT) are then modeled into B-DNA (Figure S12, Stage 3) as initial models for MD (Stage 4).The structures with * clash with neighboring base pairs in the context of duplex B-DNA and are not used for MD.

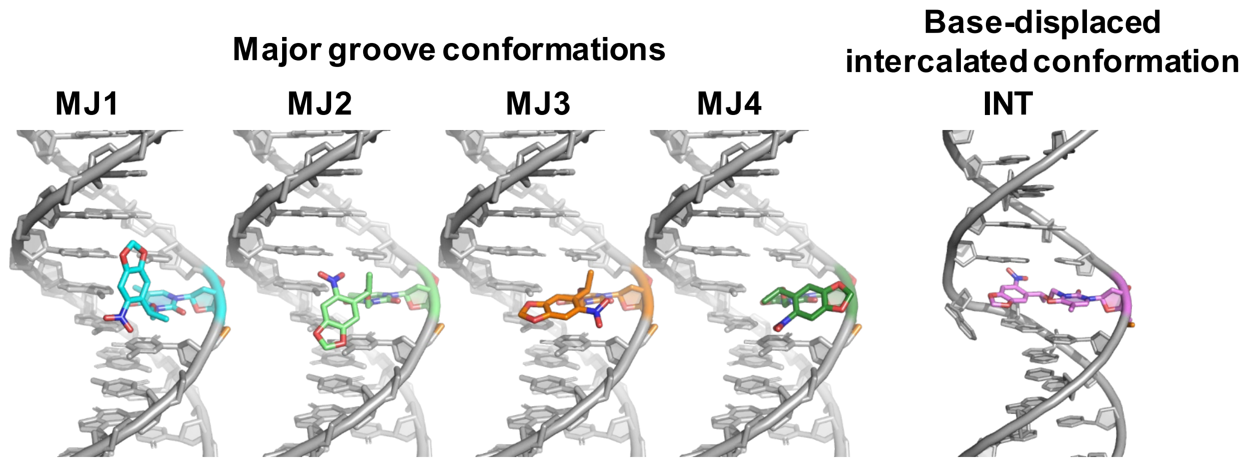


**Figure S12. Initial models of NPOM-dT-containing DNA 13mers for MD simulations** (indicated as Stage 3 in Scheme 1)**.** The NPOM-dT residues are color coded as in **Figure S11**.

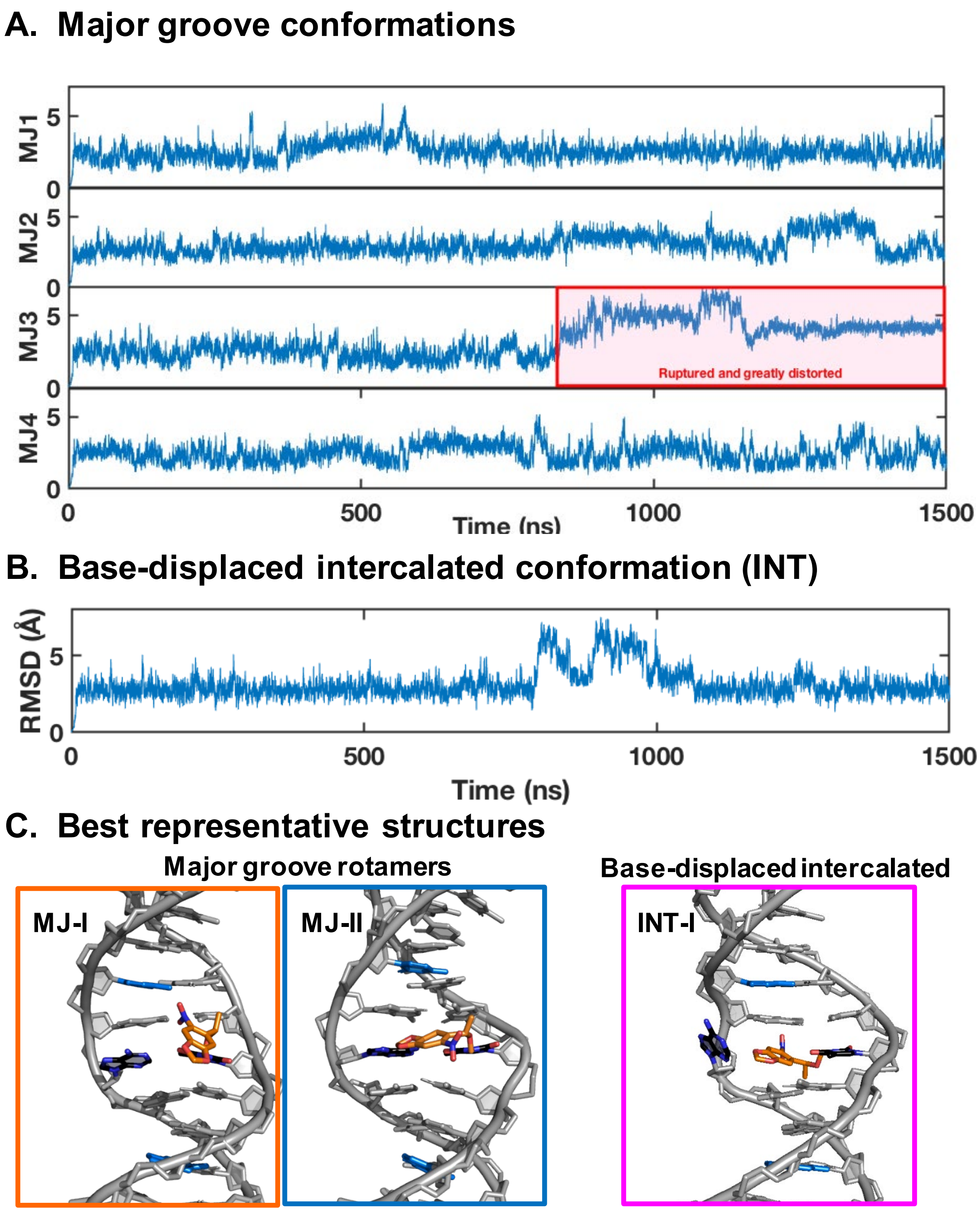


**Figure S13. RMSDs for NPOM-dT-containing 6mers in the MD simulations (A and B) and best representative structures (C).** The root-mean-squared deviation (RMSD) values are for the heavy atoms of the 6-mer sequence between the designed FRET base pair steps excluding the lesion-containing nucleotide. The best representative structures are for the two predominant major groove rotamers and one base-displaced intercalated conformation, shown in **Figure 6** in the main text (Stage 4 in Scheme 1).

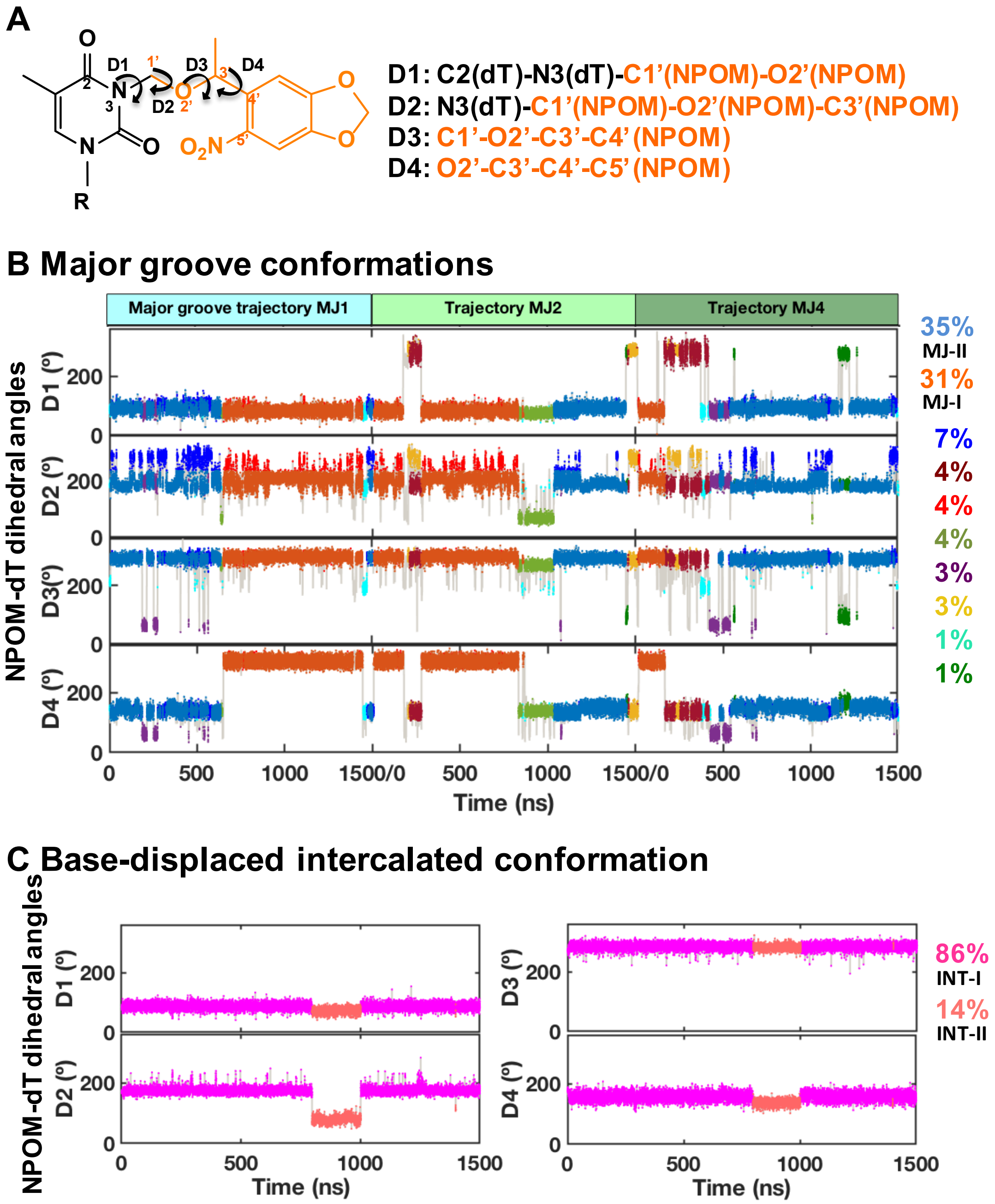


**Figure S14. Conformational clusters of the NPOM-dT based on the dihedral angles between NPOM rings and dT. (A)** The definition of dihedral angles between NPOM rings and dT. **(B)** The dihedral angle values and clusters of the NPOM-dT in the combined MD derived ensembles (MJ1, MJ2, and MJ4, Figures S12 and S13) of the major groove conformational family. **(C)** The dihedral angle values and clusters of the NPOM-dT in the MD derived ensembles of the base-displaced intercalated conformational family. Population fraction for each cluster (over 1%) is color-coded and given on the right.

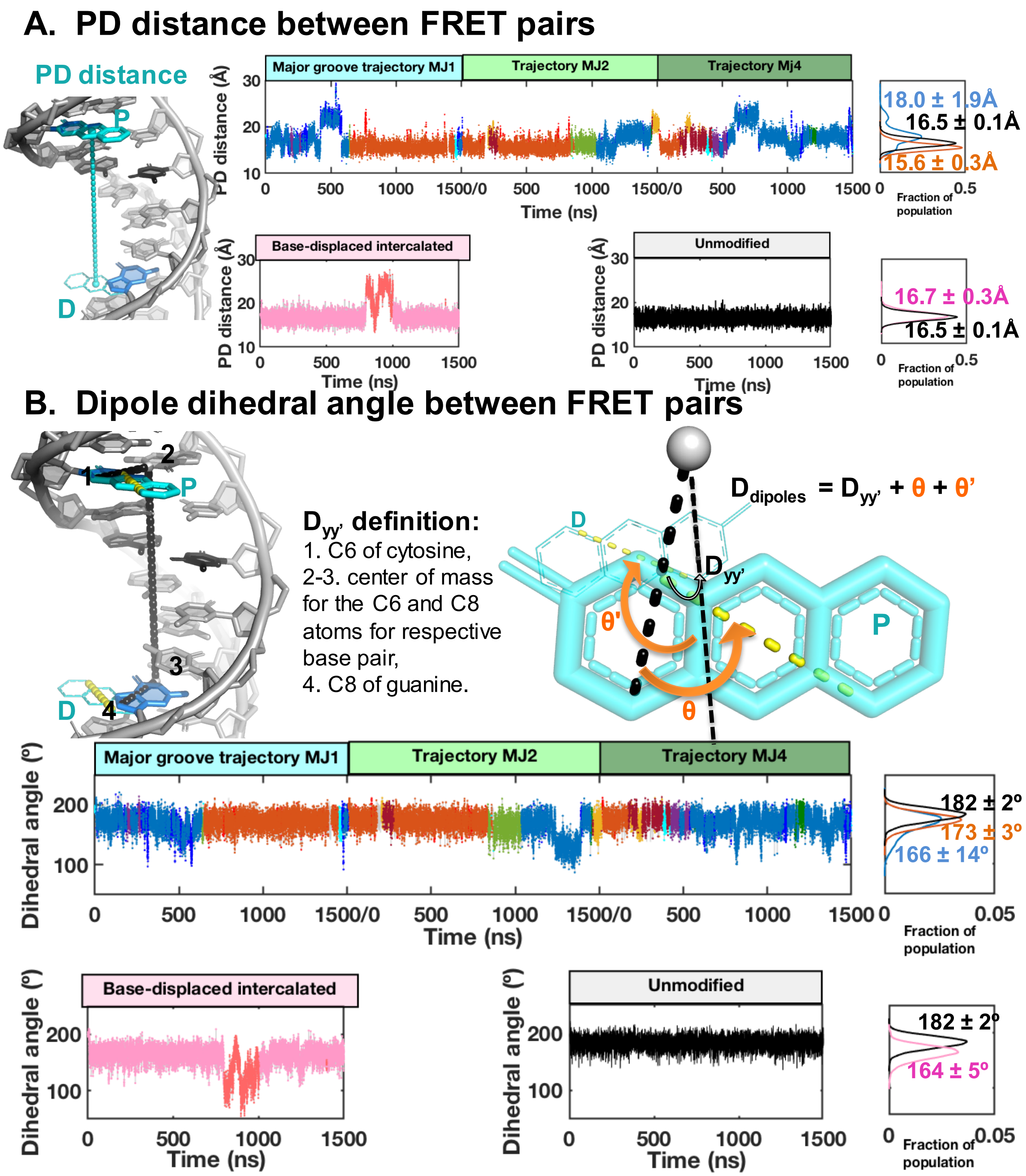


**Figure S15. The distances and the dipole dihedral angles between modeled FRET pairs.** The distances **(A)** and dipole dihedral angles **(B)** between FRET pairs are illustrated in the best-representative structure of unmodified DNA, and their values along the MD trajectories are shown. The values are color coded as in Figure S14 for each conformation with its distribution shown on the right.
